# VEGF-A/VEGFR-1: a painful astrocyte-mediated signaling blocked by the anti-VEGFR-1 mAb D16F7

**DOI:** 10.1101/2021.01.19.427244

**Authors:** Laura Micheli, Carmen Parisio, Elena Lucarini, Alessia Vona, Alessandra Toti, Alessandra Pacini, Tommaso Mello, Serena Boccella, Flavia Ricciardi, Sabatino Maione, Grazia Graziani, Pedro Miguel Lacal, Paola Failli, Carla Ghelardini, Lorenzo Di Cesare Mannelli

## Abstract

Chemotherapy-induced neuropathic pain is a clinically relevant adverse effect of several anticancer drugs leading to dose reduction or therapy discontinuation. The lack of knowledge about the mechanisms of neuropathy development and pain chronicization makes chemotherapy-induced neuropathic pain treatment an unmet medical need. In this context, the vascular endothelial growth factor A (VEGF-A) has emerged as a neurotoxicity biomarker in a model of chemotherapy-induced neuropathy, and its decrease has been related to pain relief. Aim of this study was to clarify the VEGF-A-dependent pain signaling in the CNS for individuating new targeted therapeutic approaches. In mice, the intrathecal infusion of VEGF-A induced a dose-dependent noxious hypersensitivity mediated by the VEGF receptor 1 (VEGFR-1) as demonstrated by pharmacological and genetic tools. In electrophysiological study, VEGF-A stimulated the spinal nociceptive neurons activity through VEGFR-1. In the dorsal horn of the spinal cord, VEGF-A increased in astrocytes of animals affected by neuropathy suggesting this cell population as a source of the potent pain mediator. Accordingly, the selective knockdown of astrocytic VEGF-A, by shRNAmir, blocked the development of oxaliplatin-induced neuropathic pain. Besides, the anti-VEGFR-1 mAb D16F7 (previously described as anticancer) effectively relieved neuropathic pain induced by chemotherapeutic agents. In conclusion, astrocyte-released VEGF-A is a new player in the complex neuron-glia network that oversees physiological and pathological pain and D16F7 mAb rises as a potent pain killer strategy.

## Introduction

Persistent pain induced by the neurotoxic effects of anticancer chemotherapy (hereafter referred to as chemotherapy-induced neuropathic pain, CINP) is a kind of pure neuropathic hypersensitivity. Direct nervous tissue damages and a complex maladaptive response of peripheral and central nervous systems orchestrate a painful syndrome that keeps on after therapy discontinuation and beyond cancer resolution (Di Cesare Mannelli *et al*, 2013; Miltenburg & Boogerd, 2014; Ibrahim & Ehrlich, 2020).

Studying the role of stem cells in relieving neuropathic pain, we recently showed that the plasma concentration of vascular endothelial growth factor A (VEGF-A) was enhanced in rats repeatedly treated with oxaliplatin. Stem cells were able to control pain and normalize VEGF-A suggesting a possible role of the growth factor as pain mediator (Di Cesare Mannelli *et al*, 2018).

VEGF-A is a large anti-parallel homodimeric peptide that belongs to the “Cys-loop” superfamily of proteins. It is mainly known as a pro-angiogenic factor mediating blood vessel formation, vascular permeability, endothelial cell proliferation, differentiation, leakage, migration, survival, and motility (Iyer & Acharya, 2011). Alternative splicing of the *Vegfa* gene selectively removes intron regions and joins specific combinations of exons to generate distinct VEGF-A isoforms. Differing in respect to their length, isoforms are designated as VEGF_xxx_ where xxx represents the number of amino acids present in the final protein sequence (the prototypical transcript VEGF_165_ as well as the other commonly represented VEGF_111_, VEGF_121_, VEGF_145_, VEGF_189_, and VEGF_206_; (Peach *et al*, 2018). A major site of alternative splicing occurs at exon 8, whereby proximal splicing results in the VEGF_xxx_a forms and distal splicing generates the VEGF_xxx_b isoforms containing exon 8b (Stevens & Oltean, 2018). VEGF_xxx_a isoforms (that only differ from VEGF_xxx_b in the six amino acids found at their C termini) are considered to be “pro-angiogenic”, whereas VEGF_xxx_b isoforms have been reported to have “anti-angiogenic” properties (Peach *et al*, 2018), although this description does not reflect the whole functions of VEGF-A_xxx_b’s (Ved *et al*, 2018). Interestingly, in quiescent vessels, a higher proportion of total VEGF-A is represented by VEGF_165_b (Woolard *et al*, 2004).

Despite its discovery as an angiogenic factor, from an evolutionary perspective VEGF-A emerged in the CNS of primitive organisms that lacked an established vasculature, suggesting a vessel-independent activity (Ruiz de Almodovar *et al*, 2009; Ponnambalam & Alberghina, 2011). Indeed, growing evidence indicates a diverse range of effects of VEGF-A on neural cells during development and in adulthood (Ruiz de Almodovar *et al*, 2009; Lange *et al*, 2016). It promotes CNS perfusion and induces direct neurotrophic effects in normal and pathological conditions and, as a permeability factor, VEGF-A modulates blood-brain barrier (BBB) functionality (Argaw *et al*, 2012; Licht & Keshet, 2013).

Cellular responses to VEGF-A are mainly driven by their cognate receptors, VEGFR-1 and −2, belonging to the class IV receptor tyrosine kinase family. The well-known VEGFR-2 plays essential roles in angiogenesis (Nakayama *et al*, 2013), as well as it mediates the neuroprotective effects of VEGF-A (Taiana *et al*, 2014; Verheyen *et al*, 2013). VEGFR-1 has a higher affinity for VEGF-A than VEGFR-2 but it shows decreased tyrosine kinase activity. VEGFR-1 is widely expressed also in non-endothelial cells and its soluble forms exhibit a negative modulatory activity on VEGFR-2 effects (Failla *et al*, 2018; Peach *et al*, 2018; Seki *et al*, 2018); nevertheless, its biological functions remain largely undefined.

The role of VEGF-A in pain signalling is debated as conflicting literature data suggest both algesic (Lin *et al*, 2010; Selvaraj *et al*, 2015; Hamilton *et al*, 2016; Lee *et al*, 2019) and analgesic effects (Verheyen *et al*, 2012; Hulse *et al*, 2014; Ved *et al*, 2018; Hu *et al*, 2019; Verheyen *et al*, 2013). The present work intends to dissect the pain modulatory properties of VEGF-A in the CNS in physiological and neuropathic conditions by using preclinical *in vivo* models of CINP. Moreover, the role of the different receptor subtypes in pain signalling and the potential of the VEGF-A/VEGFRs system as target for relieving pain was explored.

## Results

### Nociceptive effect of VEGFRs selective ligands infused in spinal cord

To study the spinal impact of VEGF-A signalling modulation on pain threshold in mice, we firstly evaluated the effect of the most expressed isoform VEGF_165_b. After i.t. administration, pain sensitivity was measured as latency response to a cold stimulus (Cold plate test). As shown in Fig. 1A, VEGF_165_b (3, 10 and 30 ng, in bolus in a total volume of 5 µl) dose-dependently reduced pain threshold with a long-lasting effect starting 30 min after injection that completely disappeared only after 6 h, resembling its effect observed in rats (Di Cesare Mannelli *et al*, 2018). To note, VEGF_165_a (1, 3 and 30 ng, i.t.) evoked similar dose-dependent nociceptive effects (Supplementary Fig. S1). Since VEGF_165_ isoforms may interact with both VEGFR-1 and VEGFR-2, in order to explore the implications of the receptor types in pain modulation, we also tested the effect of placental growth factor 2 (PlGF-2) and VEGF-E, which are specific VEGFR-1 and VEGFR-2 agonists, respectively (Cudmore *et al*, 2012). As shown in Fig. 1B and 1C, both PlGF-2 and VEGF-E (3, 10 and 30 ng, i.t.) significantly reduced the licking latency of animals challenged on a cold surface (Cold Plate test), even if PlGF-2 showed a profile similar to VEGF_165_b while VEGF-E exhibited a lower efficacy. Interestingly, the selective VEGFR-1 blockade by the anti-VEGFR-1 mAb D16F7 (1 μg, i.t.), in the absence of VEGF_165_b, did not significantly alter pain threshold at microgram dose (Fig. 1D). On the contrary, nanogram dose of the anti-murine VEGFR-2 mAb DC101 (1 and 6 ng, i.t.) induced hypersensitivity (Fig. 1E) and this effect was blocked by D16F7 mAb (10 and 100 ng; Fig. 1F). In this test, non-specific mouse IgG (1 µg), used as control, was inactive. These findings suggested that the nociceptive effects evoked by VEGF_165_b were the result of VEGFR-1 stimulation. Furthermore, algesic effects induced by the DC101 mAb were likely due to the antibody-dependent displacement of the endogenous VEGF-A from VEGFR-2, thus making it available for binding to VEGFR-1; this hypothesis was further demonstrated by the loss of the effect when the anti-VEGFR-1 mAb D16F7, was administered together with DC101.

**Figure 1.**
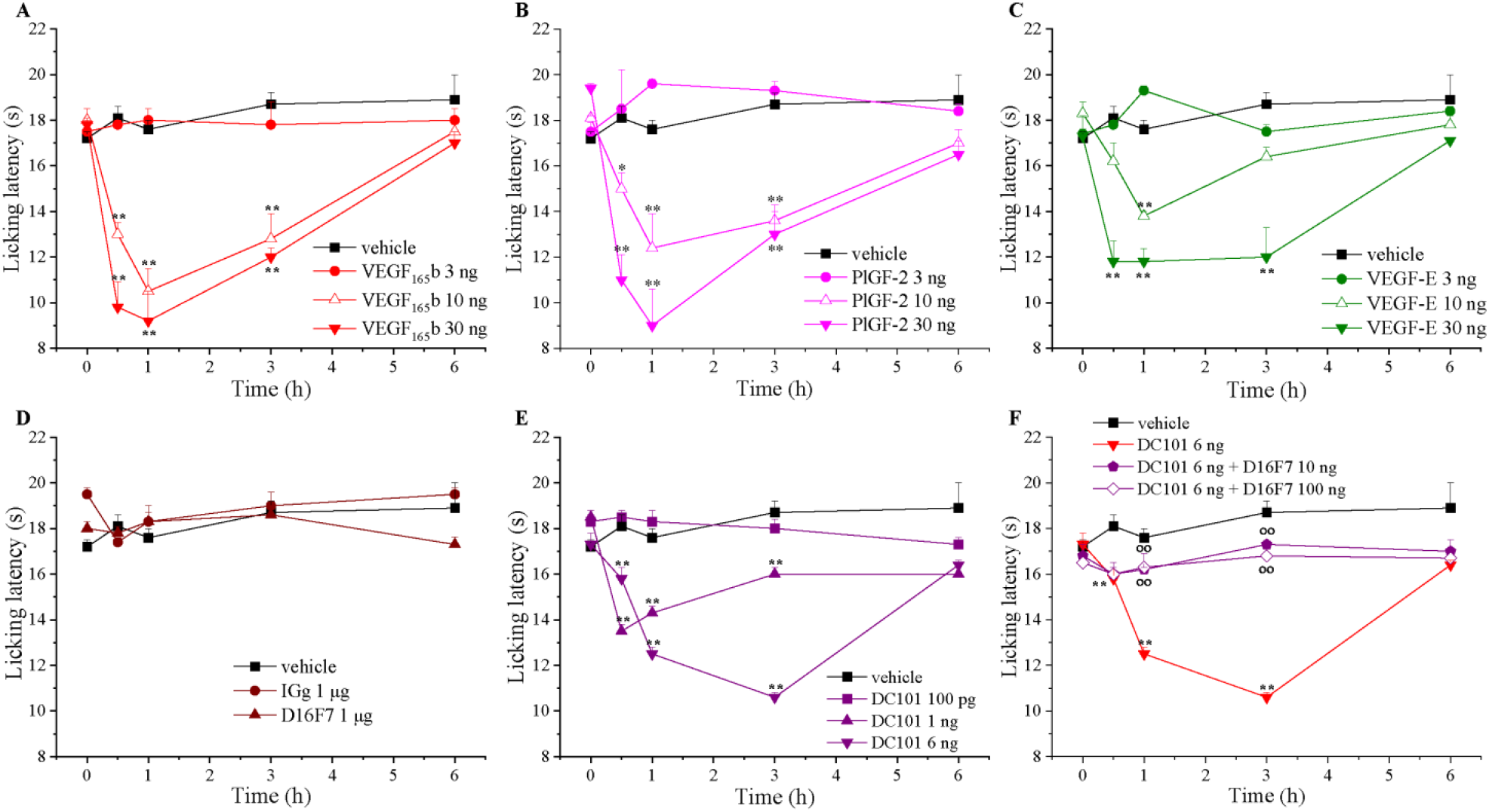
Nociceptive effect of VEGFRs selective ligands infused in spinal cord. The pain threshold was measured by the Cold plate test over time after the i.t. injection of compounds. Effect of (A) VEGF165b (n=7), (B) the selective VEGFR-1 agonist PlGF-2 (n=5), (C) the selective VEGFR-2 agonist VEGF-E (n=5), (D) the selective anti-VEGFR-1 antibody D16F7 (n=7) or a murine control IgG (n=5) and (E) the selective anti-VEGFR-2 antibody DC101 (n=5). (F) Effect of DC101 in mice pre-treated (15 min before) with D16F7. Each value represents the mean ± SEM. ^*^P<0.05 and ^**^P<0.01 vs vehicle-treated animals; °°P<0.01 vs DC101 6 ng treated animals. The analysis of variance was performed by One-way ANOVA. A Bonferroni’s significant difference procedure was used as post hoc comparison.

### Hypersensitivity-induced by VEGF-A signalling modulators is due to VEGFR-1 activation

The hypothesis that VEGFR-1 activation is required for VEGF_165_b-mediated nociception was demonstrated by crossing the combinations of receptor agonists and antagonists. Both selective agonists, PlGF-2 and VEGF-E (Cai *et al*, 2017; Park *et al*, 1994; Persico *et al*, 1999; Meyer, 1999), share the same binding sites of VEGF-A on the corresponding receptors. At variance with DC101 mAb which is a competitive antagonist of VEGF-A and VEGF-E for VEGFR-2 binding (Falcon *et al*, 2016), D16F7 mAb is a non-competitive antagonist since it interacts with VEGFR-1 at a site different from that used by the receptor ligands (Graziani *et al*, 2016; Lacal & Graziani, 2018). Consistently with our hypothesis, the algesic effects of VEGF_165_b are blocked by D16F7 mAb (Fig. 2A). A similar profile was obtained also for the VEGFR-1 ligand PlGF-2 (Fig. 2B) as well as for the VEGFR-2 ligand VEGF-E (Fig. 2C). DC101 mAb used at the highest non-algesic dose (but able to selectively block VEGFR-2) (Falcon *et al*, 2016) did not block the effect of both VEGF_165_b and PlGF-2, but further exacerbated VEGF-E hypersensitivity (Supplementary Fig. S2). These findings confirmed the pivotal role of VEGFR-1 in pain signalling which is directly activated by the selective agonist PlGF-2 or by the exogenously added (Fig. 2A) or endogenously present VEGF-A (Fig. 2C) displaced from VEGFR-2. Moreover, the selective knockdown of VEGFR-1 or VEGFR-2 by siRNA further validated the specificity of the VEGFR-1-mediated mechanism (Fig. 2D-F). The silencing of VEGFR-1 completely blocked the effects of VEGF_165_b, PlGF-2 and VEGF-E (Fig. 2E) whereas the silencing of VEGFR-2 did not alter the algesic properties of the ligands (Fig. 2F).

**Figure 2.**
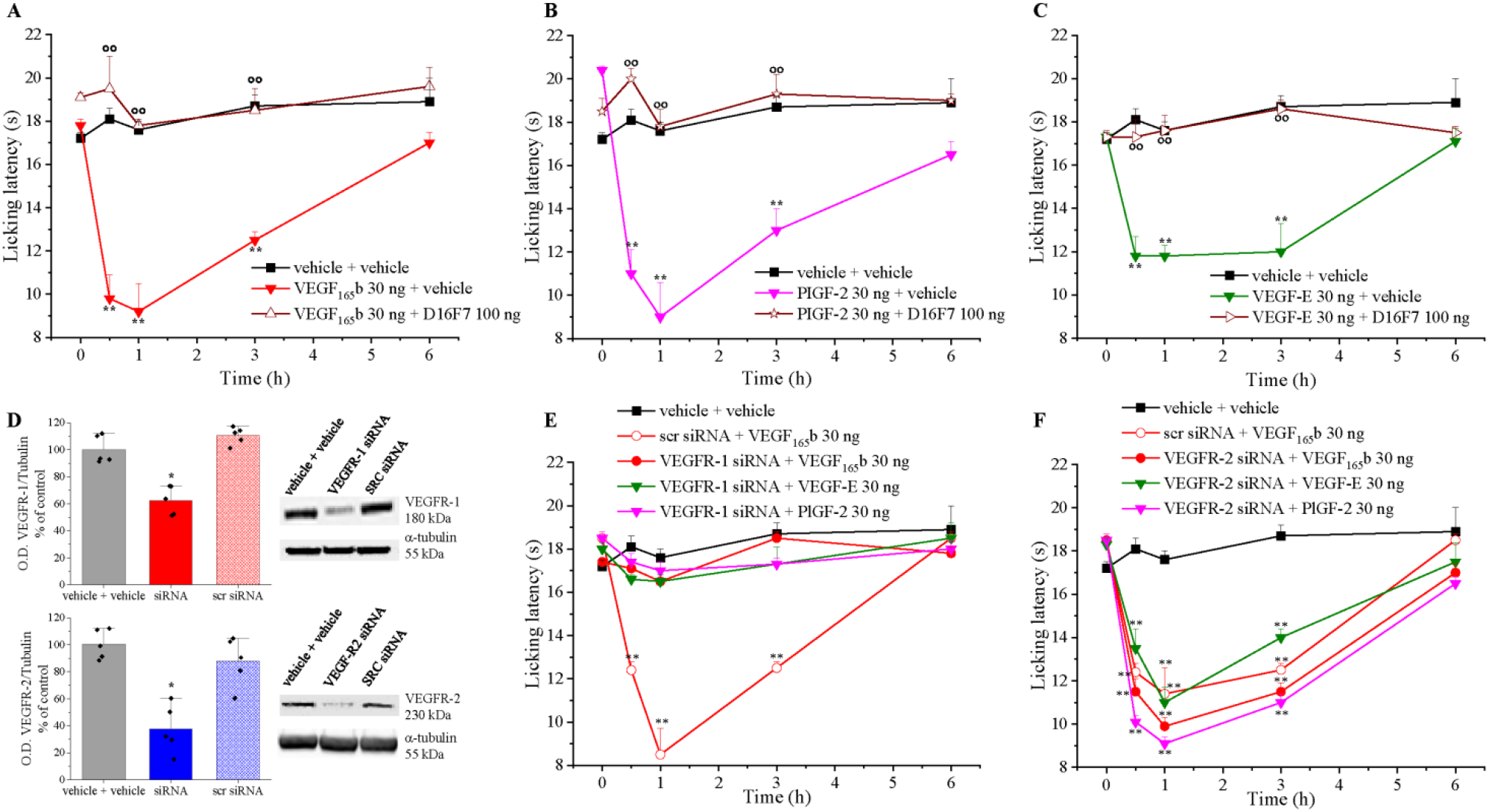
Hypersensitivity-induced by VEGF-A signalling modulators is due to its interaction with VEGFR-1. The response to a thermal stimulus (Cold plate test) was recorded after i.t. infusion of different VEGFR ligands (30 ng) preceded (15 min before) or not by the anti-VEGFR-1 mAb D16F7 (100 ng) or vehicle: (A) VEGF165b ± D16F7 (n=5), (B) PlGF-2 ± D16F7 (n=5), (C) VEGF-E ± D16F7 (n=5). (E − F) Effects of VEGFRs ligands (i.t.) in mice undergone a selective knockdown of VEGFR-1 (D, n=5) or VEGFR-2 (E, n=5) at the lumbar level of the spinal cord by siRNA. (D) Representative western blot images and densitometric analysis showing the expression of VEGFR-1 or VEGFR-2 in the lumbar section of the spinal cord after the siRNAs administration (n=5). Each value represents the mean ± SEM. ^**^P<0.01 vs vehicle + vehicle-treated animals; °°P<0.01 vs vehicle + VEGFRs ligands-treated animals. The analysis of variance was performed by One-way ANOVA. A Bonferroni’s significant difference procedure was used as post hoc comparison.

### VEGF_165_b increases the activity of spinal nociceptive neurons by VEGFR-1 activation

To investigate the effect of the spinal application of VEGF_165_b on the hyperexcitability of spinal nociceptive specific (NS) neurons, *in vivo* electrophysiological experiments were performed. The results relate to NS neurons (one cell recorded from each animal per treatment) localized at a depth of 0.7-1.0 mm from the surface of the spinal cord. This cell population was characterized by a mean rate of basal firing of 0.015 ± 0.002 spikes/sec and only cells showing this pattern were chosen for the experiment. Spontaneous and noxious-evoked (mechanical stimulation) activity of NS neurons was measured after spinal application of VEGF165b preceded or not by treatment with the anti-VEGFR-1 mAb D16F7 to investigate the involvement of VEGF-A receptor subtype. Representative ratematers of the results obtained with VEGF_165_b in the absence or presence of D16F7 mAb are shown in Fig. 3A and 3B, respectively. In mice pre-treated with vehicle (DMSO in 0.9% NaCl), VEGF_165_b (3 ng/5 µl) spinal application induced an increase in spinal electrical activity as compared to baseline levels (100%). In particular, NS neurons showed a variation of spontaneous activity compared to baseline of 217.05 ± 29.2% as well as a noxious-evoked activity with frequency of 234 ± 30.9% and duration of 316.2 ± 27.2%, starting from 25 min post VEGF_165_b (Fig. 3A, C-E). The spinal VEGF_165_b-induced hypersensitivity was mainly mediated by VEGFR-1 rather than VEGFR-2 activation. Indeed, electrophysiological recordings revealed that spinal pre-application of D16F7 mAb (100 pg/5 µl) significantly prevented the increase of spontaneous and noxious-induced activity of NS neurons resulting in a pattern similar to basal (Fig. 3B, C-E). D16F7 (100 pg/5 µl) alone was not able to affect either spontaneous or evoked activity of NS neurons (Fig. 3B). On the contrary, DC101 at 30 and 100 pg, showed a pro-nociceptive effect on NS spinal activity *per se* (Supplementary Fig. S3, representative ratematers). In fact, post-injection level of either spontaneous (187.3 ± 17.7% at 100 pg and 151.1 ± 6.9% at 30 pg) or noxious pressure-evoked firing rates (frequency: 212.6 ± 27% at 100 pg and 152.9 ± 6.9% at 30 pg; duration: 235.7 ± 25.3% at 100 pg and 119.7 ± 8.6% at 30 pg) were significantly higher respect to the baseline, in a dose-dependent manner. Overall, these results further confirmed the involvement of VEGFR-1 in VEGF-A-induced electrophysiological changes of NS.

**Figure 3.**
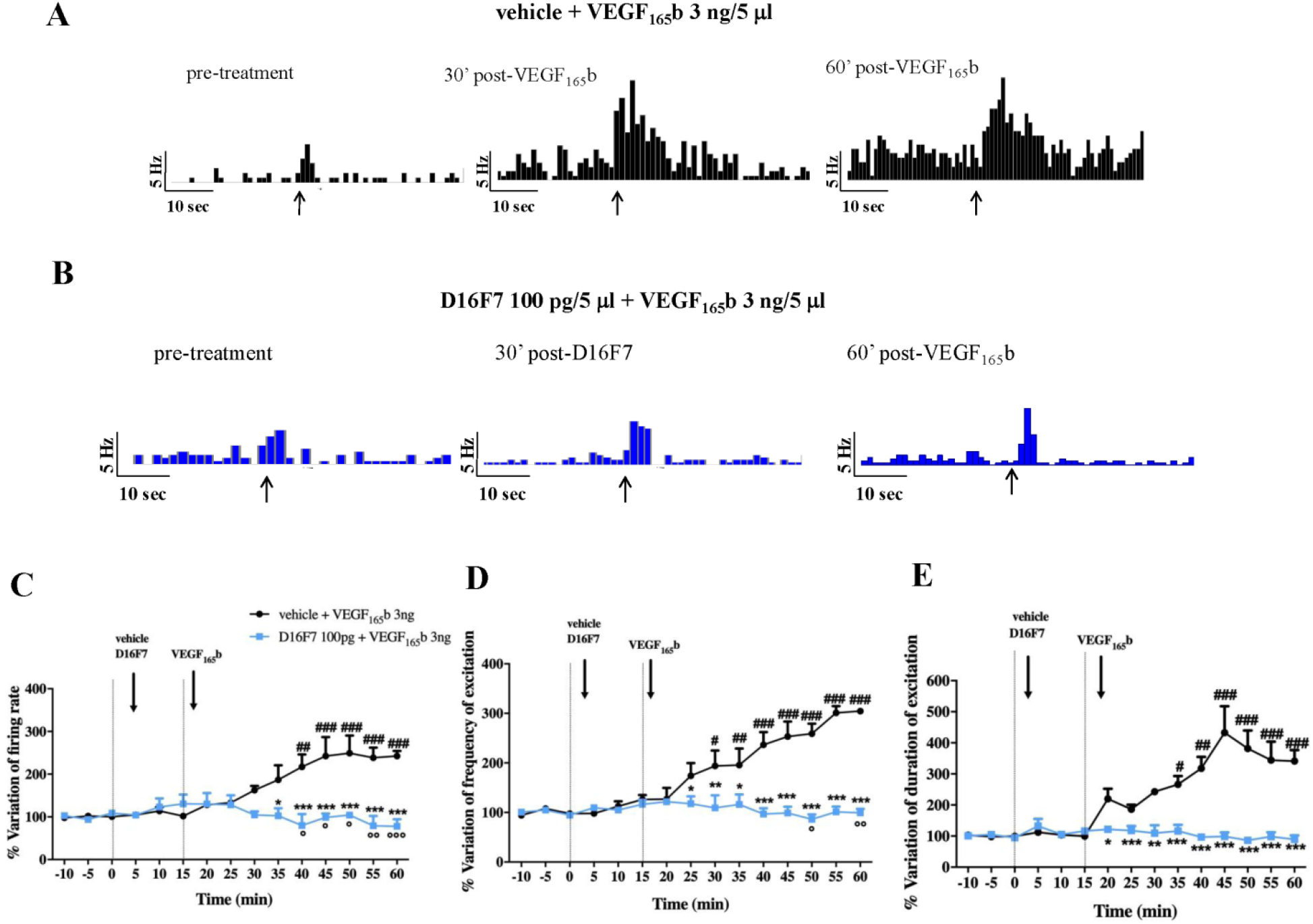
VEGF165b increases spontaneous and noxious-evoked activity of NS neurons by VEGFR-1. Representative ratematers showing spontaneous and noxious-evoked activity of NS neurons after spinal application of VEGF165b alone or in combination with D16F7 mAbs (A and B, respectively); black arrows indicate the noxious stimulation on the mouse hind-paw. Mean ± S.E.M. population data of spinal cord application of VEGF165b (3 ng/5 □l) in the presense of vehicle (DMSO in 0.9% NaCl), or D16F7 (100 pg/5 □l) on firing rate of spontaneous activity (C), frequency (D) and duration of evoked activity (E) of NS neurons in CD1 mice. Black arrows indicate vehicle, D16F7 or VEGF165b spinal application. Each point represents the mean of 5 different mice per group (one neuron recorded per each mouse). #P<0.05, ##P<0.01 and ###P<0.001 indicate statistically difference vs baseline; ^*^P<0.05, ^**^P<0.01 and ^***^P<0.001 indicate statistically difference vs vehicle + VEGF165b. One-way ANOVA followed by Dunnet’s multiple comparison post-hoc test was performed for statistical significance within groups. Two-way ANOVA followed by Bonferroni post-hoc test was used for comparison between groups.

### VEGF-A and VEGFR-1 localization in the spinal cord of naïve mice

Immunofluorescence analysis was performed in the dorsal horn of the spinal cord to study VEGF-A and VEGFR-1 expression profile in the nervous cells. VEGF-A immunoreactivity in astrocytes (as colocalization with GFAP; Fig. 4A and 4D) was significantly higher in comparison to microglia (Iba1 positive cells) and neurons (NeuN positive cells) (Fig. 4B, 4C and 4D). As expected, VEGF-A staining is strictly related to vessel structure (Fig. 4) since its expression was observed both on endothelial cells and astrocyte endfeet (Lange *et al*, 2016). To better investigate this aspect, we compared the co-localization of VEGF-A with GFAP and RECA-1, a marker of endothelial cells. As shown in Fig. 5A, it is possible to identify separate areas of VEGF-A/GFAP and VEGF-A/RECA-1 colocalization. Furthermore, VEGF-A expression in astrocytes was also confirmed by confocal microscopy. Results shown in Fig. 5B and 5C confirm the colocalization of VEGF-A with GFAP and Aquaporin 4 (AQP4, a marker of astrocytic endfeet). Indeed, the Van Steensel’s cross-correlation function (CCF) clearly shows that VEGF-A co-localizes with GFAP and AQP4 in cellular structures with an estimated diameter of 1.00 ± 0.11 µm and 1.28 ± 0.12 µm (CCF at FWHM, mean ± SD, Supplementary Figure S4C and S4F), respectively, which are compatible with the size of astrocytic processes. Collectively, these analyses demonstrate the presence of a VEGF-A pool in astrocytes.

**Figure 4.**
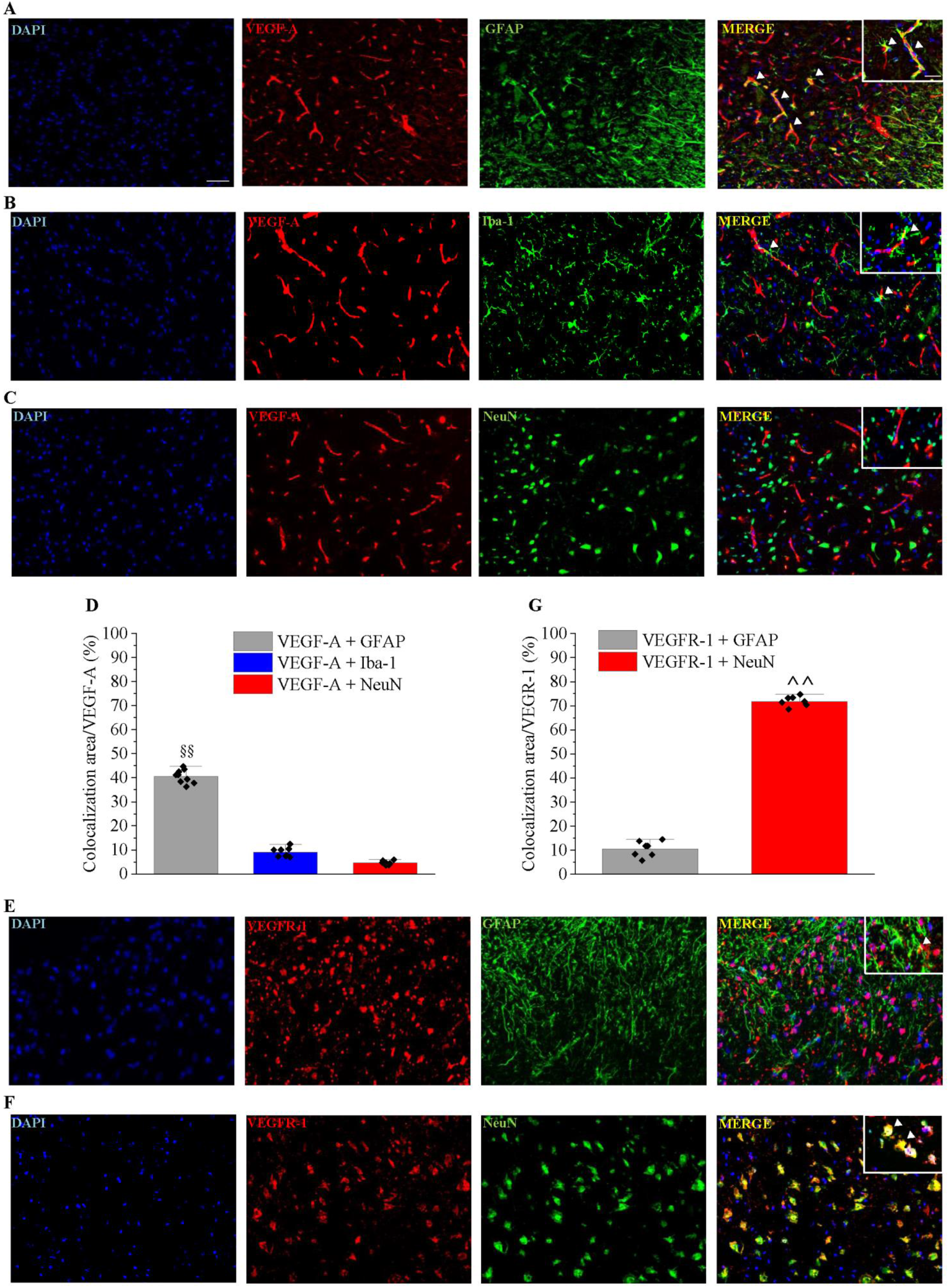
Cellular localization of VEGF-A and VEGFR-1 in the spinal cord of naïve mice. VEGF-A immunoreactivity was analysed in the spinal cord dorsal horn of naïve mice. Colocalization with GFAP-positive astrocytes (A, n=9), Iba1 positive microglia (B, n=8) and NeuN-positive neurons (C, n=7) was evaluated and quantified (D). Immunofluorescence co-staining of VEGFR-1 in the dorsal horn with GFAP (E, n=7) or NeuN (F, n=8) positive cells, and quantitative analysis (G). Scale bar: 100 μm. Each value represents the mean ± SEM. §§P<0.01 vs VEGF-A + Iba1 and VEGF-A + NeuN. ^^P<0.01 vs VEGFR1 + GFAP.

**Figure 5.**
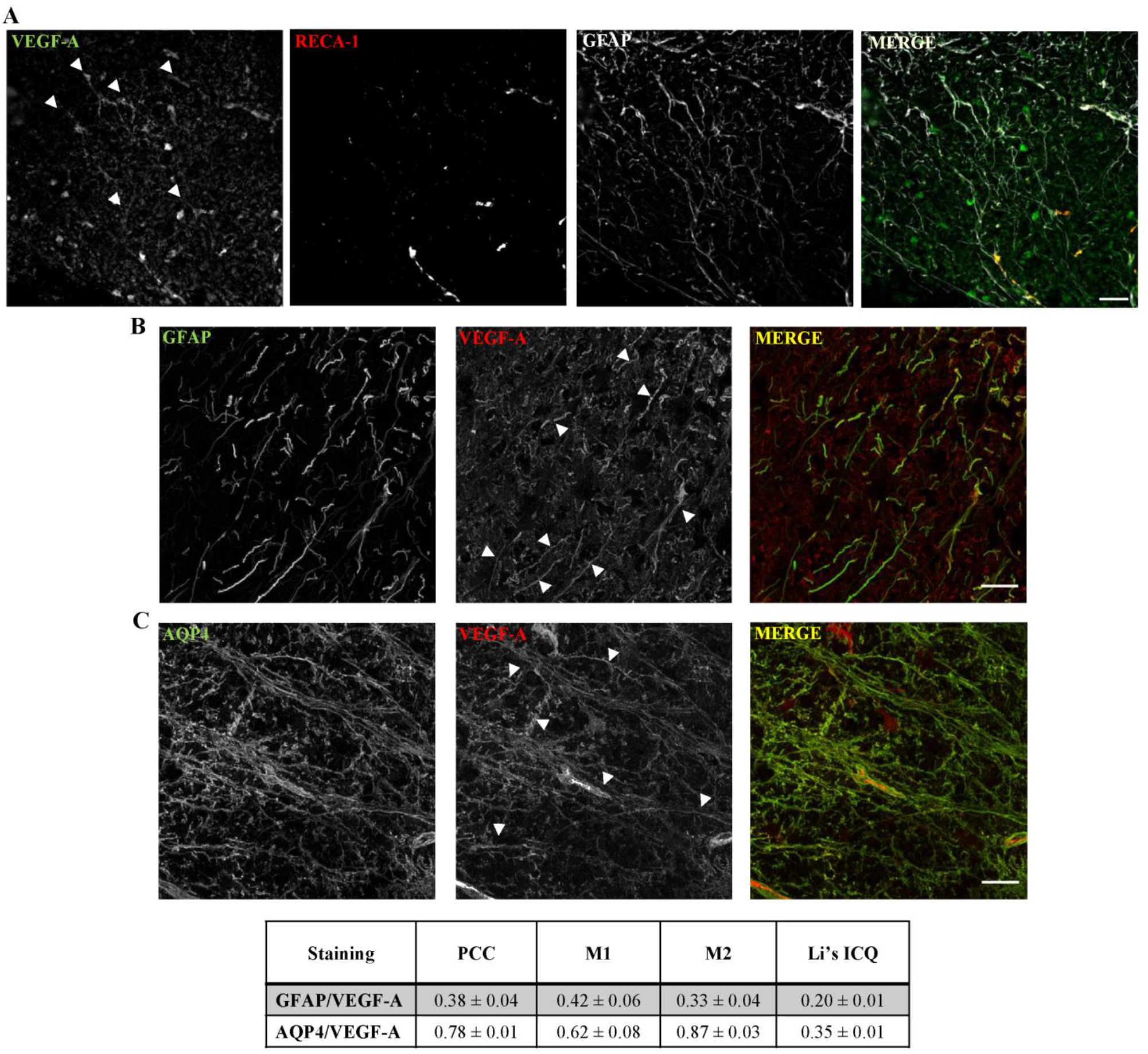
Colocalization analysis of VEGF-A and RECA-1, GFAP or AQP4 in the spinal cord of naïve mice. A) VEGF-A immunoreactivity was analysed in the spinal cord dorsal horn of naïve mice in comparison to RECA-1-positive endothelial cells and GFAP-positive astrocytes; arrows indicate the presence of VEGF-A in astrocytes; scale bar: 100 μm. B and C). Deconvolved confocal z-stacks shown as maximum intensity projection. Arrows indicate points of interest. B) Representative GFAP and VEGF-A z-stack. C) Representative GFAP and Aquaporin-4 z-stack. Table). Colocalization parameters are given as mean ± SEM (n=8), PCC= Pearson’s Correlation Coefficient; M1=Mander’s M1; M2= Mander’s M2; Li’s ICQ= Li’s Intensity Correlation Quotient. Colocalization graph are shown in Supplementary Figure S4.

As regards the expression of VEGFR-1, it was more prominent in neurons than in astrocytes (Fig. 4E, 4F and 4G).

### VEGF-A is increased in spinal astrocytes of mice with oxaliplatin-induced neuropathy

A painful neuropathy was reproduced in mice by a repeated treatment with oxaliplatin (Cavaletti *et al*, 2001; Di Cesare Mannelli *et al*, 2017). After 2 weeks of treatment, when hypersensitivity was developed, VEGF-A immunoreactivity significantly increased in dorsal horns of the spinal cord in comparison to control animals (Fig. 6A and Supplementary Fig. S5). The increment was specifically confirmed in astrocytes when colocalization of VEGF-A expression in GFAP-positive cells was measured (Fig. 6B, 6C and 6D). As regards VEGFRs, VEGFR-2 expression increased in the spinal cord of oxaliplatin-treated mice as revealed by western blot, on the contrary VEGFR-1 was unaffected by chemotherapy (Supplementary Fig. S6).

**Figure 6.**
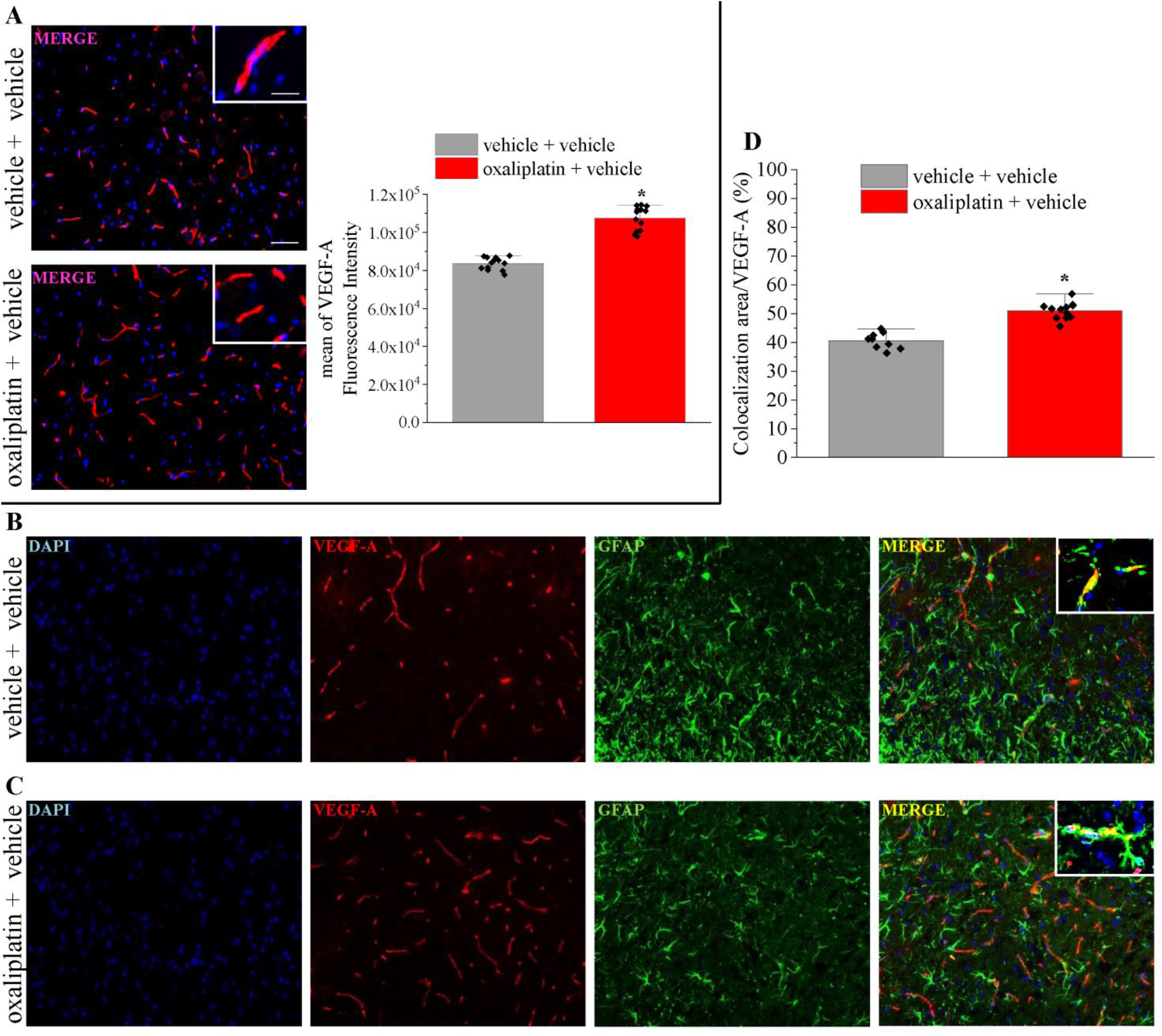
VEGF-A is increased in spinal astrocytes of mice with oxaliplatin-induced neuropathy. (A) Representative images and quantitative analysis of mean VEGF-A fluorescence intensity in the dorsal horn of oxaliplatin-treated mice in comparison to control (n=13). (B-D) Colocalization analysis of VEGF-A and GFAP in the different groups, a quantitative analysis was reported (D) (vehicle + vehicle, n=13; oxaliplatin + vehicle, n=12). Scale bar: 100 μm; insert: 50 μm. Each value represents the mean ± SEM. ^*^P<0.05 vs vehicle + vehicle group. The analysis of variance was performed by One-way ANOVA. A Bonferroni’s significant difference procedure was used as post hoc comparison.

### VEGF-A silencing in astrocytes prevents neuropathic pain

To study the influence of astrocytic VEGF-A modulation on pain signaling, we selectively silenced VEGF-A in spinal astrocytes by injecting an AAV1-GFAP-eGFP-VEGFA-shRNAmir. The vector was bilaterally injected at the lumbar and thoracic levels of the spinal cord 2 weeks before the first oxaliplatin treatment. As shown in Fig. 7A, 4 weeks after injection, the vector fluorescence colocalized with GFAP-positive cells inducing a significant decrease of VEGF-A expression (Fig. 7B). The pain threshold measurements by employing thermal (Cold plate test) and mechanical (Von Frey test) non-noxious stimuli over time showed a significant prevention of hypersensitivity development during the 2 weeks of oxaliplatin treatment in the group that received the VEGF-A specific shRNAmir in comparison to scrambled- and vehicle-treated mice (Fig. 7C and 7D). To verify the lack of neurological and motor alterations which could interfere with pain behavior recordings, VEGF-A shRNAmir and scrambled-treated mice motor functionality and exploratory activity were evaluated by the Hole board test. No alterations were highlighted with the exception of a higher exploratory activity on day 3 of oxaliplatin protocol (Supplementary Table S2).

**Figure 7.**
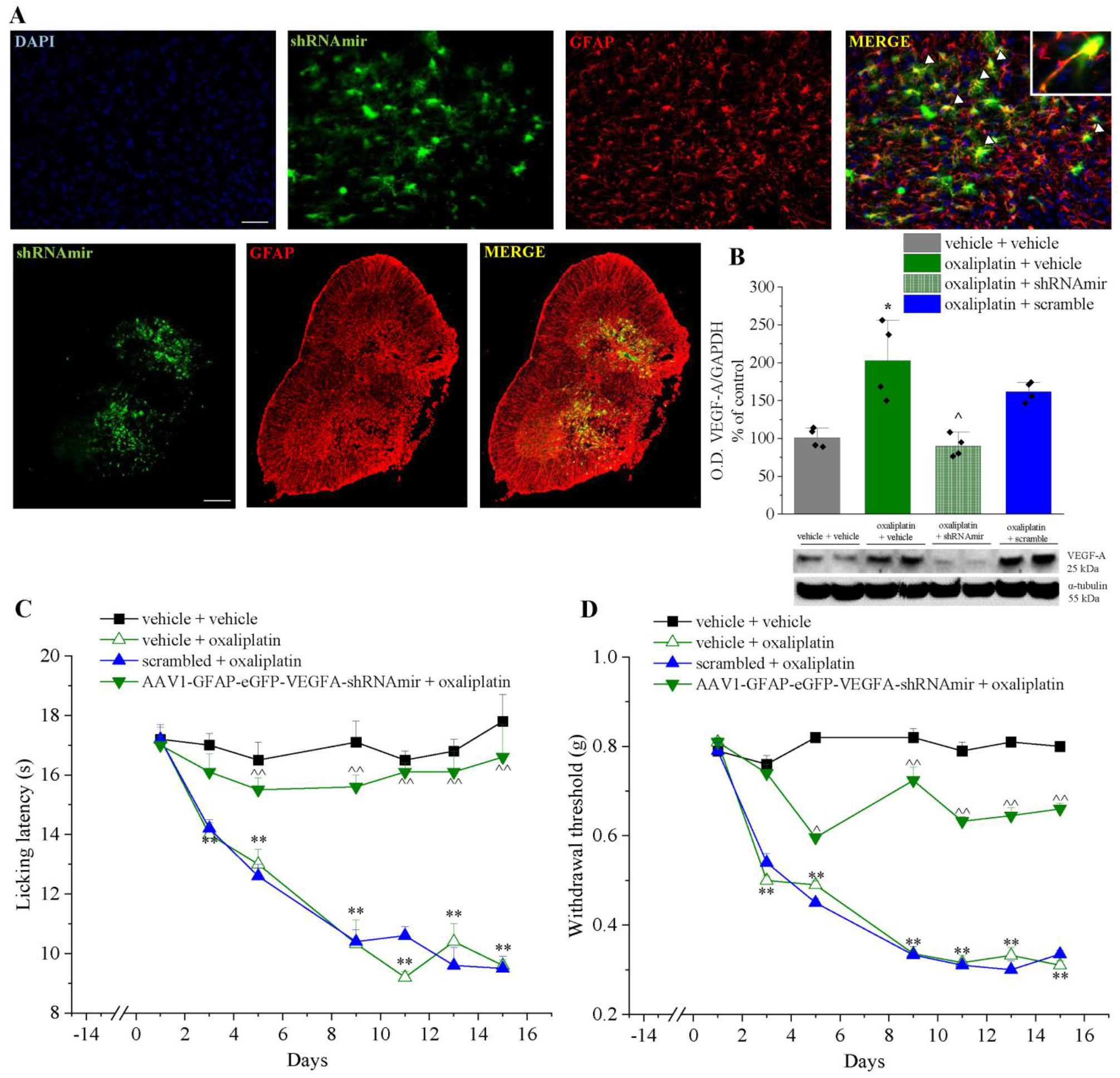
VEGF-A silencing in astrocytes prevents neuropathic pain. AAV1-GFAP-eGFP-VEGFA-shRNAmir was injected in the spinal cord to decrease VEGF-A expression in astrocytes. (A) Representative image of eGFP and GFAP fluorescence in a whole section at lumbar level, scale bar: 100 μm. Higher magnifications were reported to visualize the colocalization, scale bar: 50 μm (n=4). (B) Representative western blot images and densitometric analysis showing the expression of VEGF-A in the lumbar section of the spinal cord after the vector administration (n=4, blot of samples obtained from 2 animals of each group are shown). Pain threshold was evaluated by (C) Cold plate and (D) Paw pressure test (n=5). Each value represents the mean ± SEM. ^*^P<0.05 and ^**^P<0.01 vs vehicle + vehicle; ^P<0.05 and ^^P<0.01 vs scrambled + oxaliplatin group. The analysis of variance was performed by One-way ANOVA. A Bonferroni’s significant difference procedure was used as post hoc comparison.

### The anti VEGFR1 mAb D16F7 relieves pain in different models of CINP

To evaluate the anti-hypersensitivity potential of D16F7, we tested its activity against neuropathic pain induced by anticancer drugs. In the already described oxaliplatin model, D16F7 was infused i.t. (100 ng, 1 µg and 5 µg) showing a significant, dose-dependent, increase of the pain threshold both after thermal and mechanical non-noxious and noxious stimulation, respectively. Hypersensitivity was fully counteracted (up to control values) from 30 min to 3 h after treatment (Fig. 8A and 8B). On the contrary, the anti VEGFR-2 antibody DC101 (100 pg i.t.) was ineffective (data not shown). Interestingly, D16F7 mAb maintained its efficacy also when systemically injected by the i.p. route (1, 5, 15 and 25 mg kg^-1^). It was active starting from the dose 5 mg kg^-1^, the onset of the analgesic effect was observed at 60 min and efficacy maintained up to 120 min (Supplementary Fig. S7A and S7B). The pain relieving properties of D16F7 mAb seems to be not limited to the oxaliplatin neurotoxicity since it was also effective in mice become hypersensitive after treatment with the neurotoxic anticancer drugs vincristine and paclitaxel. In both models, D16F7 mAb (1 and 5 µg, i.t.) was active between 30 min and 3 h (Fig. 8C-F) in the Cold plate and Paw pressure tests with a particular efficacy when the pain response was evoked by thermal stimuli (Fig. 8C and 8E). In paclitaxel-treated mice 15 µg D16F7 mAb dosed i.t. was effective till to 5 h (Fig. 8E).

**Figure 8.**
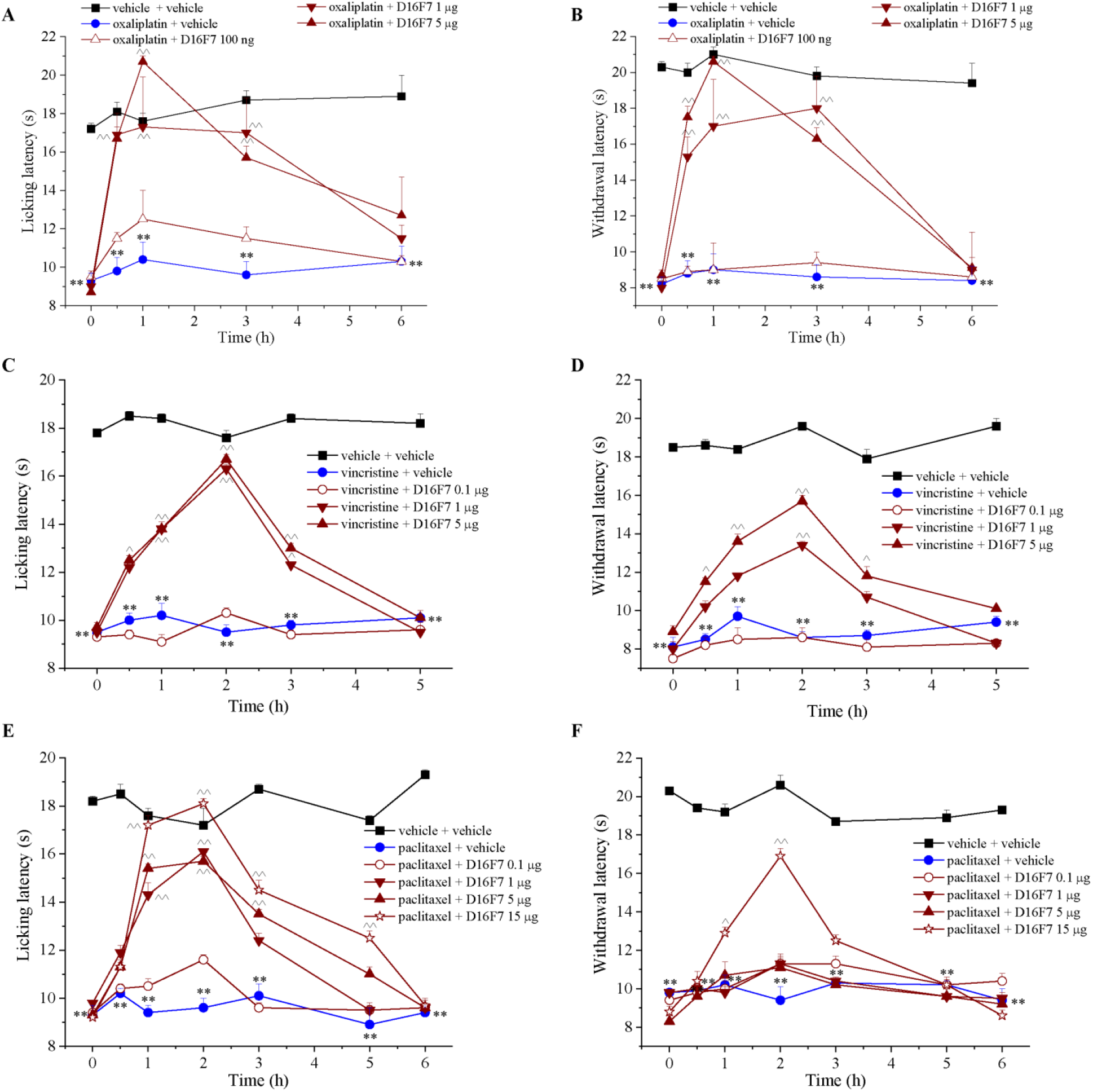
D16F7 mAb reduces pain in different models of chemotherapy-induced neuropathy. Effect of D16F7 mAb evaluated by (A) Cold plate and (B) Paw pressure tests in a mouse model of oxaliplatin-induced neuropathy after i.t. injection (A, B, n=6). (C, D) Effect of D16F7 mAb after i.t. administration in vincristine-treated mice stimulated with thermal (C) or mechanical (D) stimuli (n=6). (E, F) Effect of D16F7 after i.t. administration in paclitaxel-treated mice stimulated with thermal (E) or mechanical (F) stimuli (n=6). Each value represents the mean ± SEM. ^**^P<0.01 vs vehicle + vehicle-treated animals; ^P<0.05 and ^^P<0.01 vs chemotherapeutic drugs + vehicle-treated animals. The analysis of variance was performed by One-way ANOVA. A Bonferroni’s significant difference procedure was used as post hoc comparison.

## Discussion

Our data indicate that VEGF-A evokes pain through VEGFR-1 activation at the CNS site in physiological and pathological conditions. In particular, CINP is sustained by a spinal VEGF-A release from astrocytes that can be counteracted by the anti-VEGFR-1 mAb D16F7. Using recombinant VEGF_165_b, the most represented VEGF-A isoform, we showed an increase of the electrophysiological activity of nociceptive neurons in the spinal cord with a consequent, significant decrease of the pain threshold (hypersensitivity was similarly induced by the VEGF_165_a isoform). These data are in agreement with our previous results (Di Cesare Mannelli *et al*, 2018) and with the peripheral pro-nociceptive effect of VEGF-A demonstrated by Selvaraj and colleagues (Selvaraj *et al*, 2015) after an intraplantar injection as well as with the VEGF-A increase in synovial fluid of subjects afflicted by osteoarticular pain (Hamilton *et al*, 2016). Furthermore, in peripheral nervous system, anti-VEGF-A mAb treatment was found to alleviate neuropathic pain induced by the chronic constriction injury of the sciatic nerve (Lin *et al*, 2010). In our hands, both the selective VEGFR-1 agonist PlGF-2 and the selective VEGFR-2 agonist VEGF-E induced nociception after i.t. infusion. In addition, the selective anti-VEGFR-2 mAb DC101 induced hypersensitivity whereas the selective anti-VEGFR-1 mAb D16F7 did not. Based on these data, it could be hypothesized that VEGFR-1 selectively mediates pain, either directly stimulated by the exogenously added PlGF-2 or by the endogenously produced VEGF-A displaced from VEGFR-2 by VEGF-E or by the competitive DC101 mAb. In this context, it is worth noting that D16F7 mAb is a non-competitive inhibitor that hampers VEGFR-1 activation without affecting ligand binding (Graziani *et al*, 2016; Lacal & Graziani, 2018). The involvement of VEGFR-1 in algesia induced by VEGF-A was confirmed by the ability of D16F7 mAb to block the nociceptive effects of all the agonists as well as of DC101. Furthermore, the knockdown of VEGFR-1 prevented VEGF_165_b, PlGF-2 and VEGF-E effects, strongly indicating the pivotal role of this receptor in the spinal pain pathway. These data are in agreement with those described by Selvaraj and colleagues (Selvaraj *et al*, 2015) in the peripheral nervous system where VEGF-A induced nociceptive sensitization via VEGFR-1. Consistently with behavioural data, electrophysiological experiments revealed that VEGF_165_b spinal application, caused a strongly increase of both spontaneous and evoked activity of NS neurons in naïve animals. In particular, the increased responsiveness to mechanical noxious stimuli of NS neurons induced by VEGF_165_b spinal microinjection suggests that low doses of this compound were able to induce a central sensitisation, similarly to the neuropathic pain condition induced by nerve injury. In this context, the pre-application of D16F7 prevented the VEGF-A-induced neuronal hyperexcitability, ruling out the contribution of this receptor in VEGF-A-mediated painful effects. In the normal healthy CNS, VEGF-A regulates microvascular density, vessel permeability, and maintains endothelial cell fenestration in the choroid plexus, stimulates neural stem cell proliferation and promotes neurogenesis (Lange *et al*, 2016). In pathological conditions (besides beneficial vascular effects), it safeguards stressed neurons, induces axon extension and branching, and promotes synaptic plasticity; furthermore, VEGF-A triggers proliferation, survival and migration of astrocytes and stimulates expression of trophic factors by astrocytes and microglia (Ruiz de Almodovar *et al*, 2009; Lange *et al*, 2016). Glia cells play a crucial role in the maladaptive plasticity of the nervous system in chronic pain and, particularly, in neuropathies (Scholz & Woolf, 2007). Activated by neuronal damage or by signals from periphery, glia participate in pain development and chronicization, amplifying the excitatory synaptic microenvironment (Stockstill *et al*, 2018; Di Cesare Mannelli *et al*, 2015). Released soluble factors, like cytokines and growth factors, possess a direct nociceptive effect (Sommer *et al*, 2018). Among the latter, NGF, BDNF, GDNF, etc, seem unable to separate the neuroprotective effect from the algic one probably following the evolutionary positive alarm role of physiological pain (Alles & Smith, 2018; Nencini *et al*, 2018; Garraway & Huie, 2016). In this context, the nociceptive effect of VEGF-A is not surprising. Neuropathies induced by trauma to a peripheral nerve (Hulse *et al*, 2014) or by chemotherapy (Di Cesare Mannelli *et al*, 2018) are characterized by enhanced spinal concentration of VEGF-A. The present results show, as expected (Lange *et al*, 2016), a relevant spinal VEGF-A concentration in the vessel structure; nevertheless, the existence of an *extra*-endothelial component was verified and confirmed. In comparison to microglia and neurons, astrocytes of healthy mice showed the highest amount of VEGF-A, which was clearly distinguishable from the vascular component. The repeated treatment with oxaliplatin up to the development of painful neuropathy significantly enhanced the presence of the growth factor in astroglia. The selective VEGF-A knockdown in dorsal horn astrocytes at the lumbar and thoracic levels of the spinal cord strongly reduced oxaliplatin-dependent neuropathic pain suggesting astrocytic VEGF-A as a relevant component of the pain signalling orchestrated by glia. Furthermore, enhanced concentrations of VEGF-A can also lead to other pathological alterations related to neuropathies like an increased BBB permeability (Di Cesare Mannelli *et al*, 2014; Branca *et al*, 2018; Montague-Cardoso *et al*, 2020). As low maintenance levels of VEGF-A are necessary for the integrity of the BBB, high levels can alter permeability and compromise CNS (Argaw *et al*, 2012; Licht & Keshet, 2013). The hypoxia inducible factor-1 driven by IL-1 promotes VEGF-A release from astrocytes that induces down-regulation or loss of endothelial tight proteins claudin-5 and occludin, determining a loss of BBB function (Argaw *et al*, 2012; Chapouly *et al*, 2015) by mechanisms involving VEGFR-1 (Lee *et al*, 2019). On the other hand, the increase in VEGF-A levels in neurotoxic conditions is generally related to hypoxia, clearly demonstrated in diabetic-as well as in chemotherapy-induced neuropathies (Ved *et al*, 2018; Rojas *et al*, 2018; Di Cesare Mannelli *et al*, 2018), suggesting the need of improving vascular functions (Lange *et al*, 2016). The rescue role of VEGF-A is also based on its extra-vascular neuroprotective and neurodegenerative properties mainly due to the activation of the VEGFR-2. VEGF-A stimulates the migration and survival of Schwann cells (Schratzberger *et al*, 2000), it protects neurons against chemotherapy-induced cytotoxicity via activation of VEGFR-2 and MEK1/2 and inhibition of caspase-3 (Beazley-Long *et al*, 2013). VEGF-A-signalling through VEGFR-2 leads to the protection of dorsal root ganglion sensory neurons in models of drug (paclitaxel) or hyperglycaemia-induced neuropathies, through induction of Heat Shock Protein 90 deacetylation and increase of Bcl-2 (Verheyen *et al*, 2012, 2013). The loss of endothelial VEGFR-2 signalling leads to tissue alteration in the dorsal horn and the development of hyperalgesia whereas neuronal overexpression of VEGFR-2 in mice reduced the sensitivity to paclitaxel-induced peripheral neuropathy (Verheyen *et al*, 2012). This outcome seems to be related to neuroprotective effects and, accordingly, we showed an increase of VEGFR-2 spinal expression in oxaliplatin-treated mice that could be considered an adaptive response to the damage. On the contrary, the acute stimulation of VEGFR-2 does not directly interfere with pain.

Our data show that VEGF-A induces pain by selectively activating the VEGFR-1, which is expressed on spinal sensory neurons. A dichotomy between the pro-algesic VEGFR-1-signaling and the protective VEGFR-2-signaling is suggested, offering the possibility to relieve pain through a target that conserves the neuroprotective effects of the endogenous VEGF-A. In this view, the selective anti-VEGFR-1 mAb D16F7 induced a potent pain-relieving effect against nociception triggered by VEGF-A or PlGF-2 as well as against neuropathic pain evoked by the neurotoxic adverse reactions of different anticancer drugs like oxaliplatin, paclitaxel and vincristine. In addition, the pain-relieving effect of D16F7 was demonstrated after local (i.t.) and systemic (i.p.) administration. D16F7 mAb is able to inhibit VEGFR-1 homodimerization, auto-phosphorylation and downstream signal transduction (Graziani *et al*, 2016; Atzori *et al*, 2017, 2018) and down-modulates membrane receptor signaling without affecting VEGF-A or PlGF binding (Graziani *et al*, 2016; Lacal & Graziani, 2018). Indeed, D16F7 mAb interacts with a receptor site corresponding to amino acids 149-161 of human VEGFR-1, which is different from that involved in VEGF-A or PlGF binding (Davis-Smyth *et al*, 1998; Christinger *et al*, 2004; Graziani *et al*, 2016).

CINP is one of the most common adverse events of several first-line chemotherapeutic agents, affecting several million patients worldwide each year and reducing the benefits of effective anticancer therapies in the long-term outcome. It is not possible to predict which patients will develop symptoms and when during the chemotherapy course. Moreover, pain and sensory abnormalities may persist for months, or even years after the cessation of chemotherapy (Argyriou *et al*, 2012; Kolb *et al*, 2016). The management of chemotherapy-induced neuropathy is a significant challenge and there are no drugs approved to prevent or alleviate CINP (Ibrahim & Ehrlich, 2020). In this scenario, VEGF-A is candidate to be a possible plasmatic biomarker (Di Cesare Mannelli *et al*, 2018) strictly related to pain and the selective blockade of VEGFR-1 may offer the safety qualities requested for the pharmacological treatment of cancer patients in the presence of a possible co-treatment with chemotherapy. In fact, VEGFR-1 is mostly involved in pathological processes rather than in physiological conditions (reviewed in Lacal and Graziani) (Lacal & Graziani, 2018) and in preclinical *in vivo* studies repeated dosing schedules did not cause significant adverse effects, both as single agent or in combination with immune checkpoint inhibitors (Graziani *et al*, 2016; Atzori *et al*, 2018; Ceci *et al*, 2020; Lacal, Pedro Miguel, Atzori, MG; Ruffini, F; Scimeca, M; Bonanno, E; Cicconi, R; Mattei, M; Bernardini, R; D’Atri, S; Tentori, L; Graziani, G, 2020). Interestingly, D16F7 mAb has shown activity in orthotopic *in vivo* models of two highly aggressive tumors: melanoma and glioblastoma (Graziani *et al*, 2016; Atzori *et al*, 2017, 2018). D16F7 efficacy against glioblastoma indicates that therapeutic concentrations of the mAb are reached at the CNS level after systemic administration (Atzori *et al*, 2018). Since VEGFR-1 is expressed not only in tumor cells but also in cell subsets of tumor microenvironment, its blockade by D16F7 results in: a) inhibition of tumor-associated angiogenesis; b) reduction of myeloid progenitor mobilization and tumor infiltration by M2 macrophages/microglia; c) inhibition of invasiveness and vasculogenic mimicry of VEGFR-1 positive tumor cells (Graziani *et al*, 2016; Atzori *et al*, 2018; Lacal, Pedro Miguel, Atzori, MG; Ruffini, F; Scimeca, M; Bonanno, E; Cicconi, R; Mattei, M; Bernardini, R; D’Atri, S; Tentori, L; Graziani, G, 2020).

In conclusion, VEGF-A is a pro-nociceptive molecule that in the CNS activates neuronal firing and induces pain by VEGFR-1 stimulation. Actually, VEGF-A increases during chemotherapy-induced neuropathy, and its release from astrocytes plays a decisive role in CINP development. Moreover, the anti-VEGFR-1 mAb D16F7 is suggested as a promising candidate for the treatment of CINP, adding this property to its previously described anti-tumour efficacy. The results of this proof of concept study encourage further investigation on the most effective therapeutic schedule for a long-term pain control.

## Materials and Methods

### Study approval

All animal manipulations were carried out according to the Directive 2010/63/EU of the European parliament and of the European Union council (22 September 2010) on the protection of animals used for scientific purposes and with IASP. The ethical policy of the University of Florence complies with the Guide for the Care and Use of Laboratory Animals of the US National Institutes of Health (NIH Publication No. 85-23, revised 1996; University of Florence assurance number: A5278-01). Formal approval to conduct the experiments described was obtained from the Italian Ministry of Health (No. 171/2018-PR) and from the Animal Subjects Review Board of the University of Florence and from the Animal Ethics Committee of University of Campania of Naples. Experiments involving animals have been reported according to ARRIVE guidelines (McGrath & Lilley, 2015). All efforts were made to minimize animal suffering and to reduce the number of animals used.

### Animals

Eight week old male CD-1 mice (Envigo, Varese, Italy) weighing 20-25 g at the beginning of the experimental procedure were used. Animals were housed in the Centro Stabulazione Animali da Laboratorio (University of Florence, Italy) and in Stabulario Centralizzato di Ateneo (University of Campania “Luigi Vanvitelli”, Naples, Italy) and used at least 1 week after their arrival. Ten mice were housed per cage (size 26 cm x 41 cm); animals were fed a standard laboratory diet and tap water ad libitum and kept at 23 ± 1 °C with a 12 h light/dark cycle (light at 7 am).

### Treatments

VEGF_165_b (3 – 30 ng; 5 µl; cat. #3045-VE-025, R&D System, USA), PlGF-2 (3 – 30 ng; 5 µl; cat.465-PL/CF, R&D System, USA), VEGF-E (3 – 30 ng; 5 µl; cat. #CYT-263, Prospec, Israel), D16F7 (10 − 100 ng, 5 µg; 5 µl) (Graziani *et al*., 2016) and DC101 (100 pg, 1 – 6 ng; 5 µl; catalog. #BE0060 BioCell, Boston, MA, USA) or vehicle (0.9% NaCl 5 µl) were injected i.t in conscious mice as previously described (Hylden & Wilcox, 1980). Briefly, a 25-µl Hamilton syringe connected to a 30-gauge needle was intervertebral inserted between the L4 and L5 region, and advanced 6 mm into the lumbar enlargement of the spinal cord. Behavioural measurements were performed before and 30 min, 1 h, 3 h and 6 h after the administration of compounds. DC101 or D16F7 were injected 15 min before the VEGFR-1/2 agonists when administered in the co-treatment experiments.

The scrambled siRNA or the specific VEGFR siRNA (VEGFR-1-VEGFR-2 siRNA, Ambion Life Technologies, Monza, Italy) were intrathecally (i.t.) injected twice spaced 24 h apart (3.3 µg/5 μl per mouse) at the lumbar level of the mice spinal cord. On the third day, behavioural measurements were conducted after VEGFRs agonists administration. Mice were sacrificed between the 4^th^ and the 5^th^ days for the western blot analysis. The target sequences of the anti-mouse VEGFRs siRNAs were: VEGFR-1, sense strand 5’-GCAUCUAUAAGGCAGCGGAtt-3’ and antisense strand UCCGCUGCCUUAUAGAUGCtc-3’; VEGFR-2, sense strand 5’-CCCGUAUGCUUGUAAAGAAtt-3’ and antisense strand 5’-UUCUUUACAAGCAUACGGGct-3’.

### AAV virus infection

An AAV1-GFAP-eGFP-mVEGFA-shRNAmir (1.6 × 1013 GC/ml, Vector Biosystem Inc, Malvern, PA, USA) or scrambled were used. Mice were deeply anaesthetized by intraperitoneal (i.p.) injection of ketamine (100 mg kg^-1^) (Ketavet, MSD Animal Health, Milan, Italy) and xylazine (10 mg kg^-1^) (Rompum, 20 mg/ml, Bayer, Milan, Italy) and then were placed in a stereotaxic frame using the mouse spinal adaptor (Stoelting, Wood Dale, IL, USA). The skin was incised at Th12–L5 and the mouse muscles around the left side of the interspace between Th12 − L1 and L4 - L5 vertebrae were removed and the dura mater and the arachnoid membrane were carefully incised using the tip of a 30G needle to make a small window to allow vector infusion. Intraspinal injections were done using a 5-µl Hamilton syringe connected to a 34G needle. The needle was placed 0.5 mm lateral to the spinal midline at a depth of 0.4 mm from the dorsal surface of the spinal cord and 1 µl of vector or scrambled was bilaterally injected at 0.25 µl/min with a digital microinjector (Stoelting). The needle was left on place for another 3 min to prevent backflow. The surgical site was then sutured with 3-0 silk and mice were kept on a heating pad until recovery.

### CINP in vivo models

Mice treated with oxaliplatin (Carbosynth, Pangbourne, UK; 2.4 mg kg^-1^) were administered i.p. for two weeks (Cavaletti *et al*, 2001; Di Cesare Mannelli *et al*, 2017). Oxaliplatin was dissolved in a 5% glucose solution. Control animals received an equivalent volume of vehicle. Behavioural tests were performed on day 15 for the acute treatments. In mice injected spinally with the viral vector or with the scrambled, oxaliplatin was administered for two weeks (10 total injections) starting 14 days after surgery for the viral vector administration. Control animals received an equivalent volume of vehicle. Behavioural measurements were performed on days 3, 5, 9, 11, 13 and 15.

Mice treated with paclitaxel (Carbosynth; 2.0 mg kg^-1^) were injected i.p. on four alternate days (days 1, 3, 5 and 8) (Polomano *et al*, 2001; Di Cesare Mannelli *et al*, 2017). Paclitaxel was dissolved in a mixture of 10% saline solution and Cremophor EL, a derivative of castor oil and ethylene oxide that is clinically used as paclitaxel vehicle. Control animals received an equivalent volume of vehicle. Behavioural measurements started on day 10.

Mice treated with vincristine (Carbosynth; 0.1 mg kg^-1^) were injected i.p. for five consecutive days (Weng *et al*, 2003). Vincristine was dissolved in saline solution and control animals received an equivalent volume of vehicle. Behavioural measurements started on day 8.

### Assessment of mechanical hyperalgesia (Paw pressure test)

Mechanical hyperalgesia was determined by measuring the latency in seconds to withdraw the paw away from a constant mechanical pressure exerted onto the dorsal surface (Russo *et al*, 2012; Lucarini *et al*, 2019). A 15 g calibrated glass cylindrical rod (10 mm diameter) chamfered to a conical point (3 mm diameter) was used to exert the mechanical force. The weight was suspended vertically between two rings attached to a stand and was free to move vertically. A single measure was made per animal. A cut-off time of 40 s was used.

### Assessment of thermal allodynia (Cold plate test)

Thermal allodynia was assessed using the Cold-plate test. With minimal animal-handler interaction, mice were taken from home-cages, and placed onto the surface of the cold-plate (Ugo Basile, Varese, Italy) maintained at a constant temperature of 4°C ± 1°C. Ambulation was restricted by a cylindrical plexiglas chamber (diameter: 10 cm, height: 15 cm), with open top. A timer controlled by foot peddle began timing response latency from the moment the mouse was placed onto the cold-plate. Pain-related behaviour (licking of the hind paw) was observed, and the time (seconds) of the first sign was recorded. The cut-off time of the latency of paw lifting or licking was set at 30 s (Baptista-de-Souza *et al*, 2014).

### Assessment of mechanical allodynia (Von Frey test)

Mechanical allodynia was measured with the dynamic plantar aesthesiometer (von Frey instrument) (Ugo Basile) as described by Di Cesare Mannelli and colleagues (Di Cesare Mannelli *et al*, 2017) with minor modifications. Briefly, the mice were placed in individual Plexiglas cubicles (8.5 × 3.4 × 3.4 cm) on a wire mesh platform. After approximately 30 min accommodation period, during which exploratory and grooming activity ended, the mechanical paw withdrawal threshold was measured as the hind paw withdrawal responded to von Frey hair stimulation. The mechanical stimulus was delivered to the plantar surface of the hind paw of the mouse from below the floor of the test chamber by an automated testing device. A steel rod (2 mm) was pushed with electronic ascending force (0–5 g in 35 s). When the animal withdrew its hind paw, the mechanical stimulus was automatically withdrawn, and the force recorded to the nearest 0.1 g. Nociceptive response for mechanical sensitivity was expressed as mechanical withdrawal threshold in grams. The mean was calculated from six consecutive trials and averaged for each group of mice.

### Assessment of locomotor activity (Hole-Board test)

The locomotor activity was evaluated by using the hole-board test. The apparatus consisted of a 40 cm square plane with 16 flush mounted cylindrical holes (3 cm diameter) distributed 4×4 in an equidistant, grid-like manner. Mice were placed on the centre of the board one by one and allowed to move about freely for a period of 5 min each. Two photobeams, crossing the plane from mid-point to mid-point of opposite sides, thus dividing the plane into 4 equal quadrants, automatically signalled the movement of the animal (counts in 5 min) on the surface of the plane (locomotor activity). Miniature photoelectric cells, in each of the 16 holes, recorded (counts in 5 min) the exploration of the holes (exploratory activity) by the mice (Ghelardini *et al*, 2002).

### Electrophysiological recordings of nociceptive specific (NS) neurons

On the day of electrophysiological recordings, mice were initially anesthetized with tribromoethanol (Avertin, Winthrop laboratories, New York, NY, USA; 1.25%). After tracheal cannulation, a catheter was placed into the right external jugular vein, to allow continuous infusion of propofol (5–10 mg/kg/h, i.v.) and spinal cord segments L4-L6 were exposed by laminectomy, near the dorsal root entry zone, up to a depth of 1 mm (McGaraughty *et al*, 2010). An elliptic rubber ring (about 3×5 mm), sealed with silicone gel onto the surface of the cord, was used for topical spinal drug application and to gain access to spinal neurons. Animals were fixed in a stereotaxic apparatus (David Kopf Instruments, Tujunga, CA, USA) through clamps attached to the vertebral processes. Single unit extracellular activity of dorsal horn NS neurons was performed by using a glass-insulated tungsten filament electrode (3–5 MΩ) (FHC Frederick Haer & Co., ME, USA). Spinal neurons were defined as NS neurons, when they were responding only to high intensity (noxious) stimulation (Telleria-Diaz *et al*, 2010). In particular, to confirm NS response patterns, each neuron was characterized by applying a mechanical stimulation to the ipsilateral hind paw using a von Frey filament with 97.8 mN bending force (noxious stimulation) for 2 s until it buckled slightly (Boccella *et al*, 2015; Simone *et al*, 2008). Only neurons that specifically responded to noxious hind paw stimulation were considered for recordings. The recorded signals were visualized into a window discriminator, whose output was processed by an interface CED 1401 (Cambridge Electronic Design Ltd., UK) connected to iOS 5 PC. Spike2 software (CED, version 5) was used to create peristimulus rate histograms on-line and to store and analyze digital records of single unit activity off-line. The spontaneous and noxious-evoked neuronal activity was expressed as spikes/sec (Hz) and the effect of drugs was analyzed as % variation of firing rate, frequency and duration of excitation. After recording a stable basal activity (15 min), topical spinal application of vehicle or drugs was performed, and each extracellular recording was monitored until 45-60 min post-injection. In particular, groups of animals were divided as following: 1) VEGF_165_b (3 ng/5 µl, pro-nociceptive dose on NS neurons), 2) VEGF_165_b + DC101 (10 pg/5 µl, the highest non pro-nociceptive dose) and 3) VEGF_165_b + D16F7 (100 pg/5 µl). At the end of the experiment, animals were killed with a lethal dose of urethane.

### Western blot analysis

The lumbar spinal cord of mice was explanted and immediately frozen with liquid nitrogen. The frozen tissues were homogenized with lysis buffer containing 50 mM Tris-HCl pH 8.0, 150 mM NaCl, 1 mM EDTA, 0.5% Triton X-100 and complete protease inhibitors (Roche, Milan, Italy). The suspensions were sonicated on ice using three high intensity 10s bursts with a cooling period of 10s each burst and centrifuged at 13.000xg for 10 min at 4°C. Protein concentrations were quantified by bicinchoninic acid test. Fifty μg of tissue homogenate were resolved with prefabricated polyacrylamide gel (BOLT 4-12% Bis-Tris Plus gel; Thermo Fisher Scientific, Monza, Italy) before electrophoretic transfer to nitrocellulose membranes (Bio-Rad, Milan, Italy). The membranes were blocked with 1% BSA and 5% fat-free powdered milk in PBS containing 0.1% Tween 20 (PBST) and then probed overnight at 4°C with primary antibodies specific for VEGFR-1, VEGFR-2, VEGF-A, GAPDH or α-Tubulin (Supplementary Table S1). The membranes were then incubated for 1 h in PBST containing the appropriate secondary anti-rabbit or anti-mouse antibody (Supplementary Table S1). ECL (Enhanced Chemiluminescence Pierce, Rockford, IL, USA) was used to visualize peroxidase-coated bands. Densitometric analysis was performed using the “ImageJ” analysis software (ImageJ, NIH, Bethesda, MD, USA). Normalization for α-tubulin or GAPDH content was performed. The values were reported as percentages of controls arbitrarily set at 100%.

### Immunofluorescence staining and confocal imaging

Mice were sacrificed, the L4/L5 segments of the spinal cord were exposed from the lumbovertebral column via laminectomy and identified by tracing the dorsal roots from their respective DRG. Formalin-fixed (and no-fixed, used for VEGFR-1 primary antibody) cryostat sections (7 μm) were washed 3x phosphate-buffered saline (PBS) and then were incubated, at room temperature for 1 h, in blocking solution (PBS, 0.3% Triton X-100, 5% albumin bovine serum; PBST). The sections were subsequently incubated with primary antibody, anti-VEGFR-1, anti-VEGF-A, or anti-AQP-4, overnight at 4°C (Supplementary Table S1). The following day, slides were washed 3× with PBS and then sections were incubated in the dark with secondary antibody, goat anti-rabbit or anti-mouse IgG labeled with Alexa Fluor 568, in PBST at room temperature for 2 h. After 3× PBS 0.3% Triton X-100 wash for 10 min, the sections were incubated with DAPI, a nuclei-marker, at room temperature for 5 min and then the slides were mounted using Fluoromount(tm) (Life Technologies-Thermo scientific, Rockford, IL, USA) as a mounting medium.

For double immunofluorescence, on the first day, anti-Iba1 was added and the slides incubated overnight at 4°C conditions. While, the sections to be labelled for GFAP or NeuN were incubated the second day for 2 h in the dark with mouse anti-GFAP Alexa Fluor 488-conjugated or mouse anti-NeuN Alexa Fluor 488-conjugated antibodies (Supplementary Table S1). For triple immunofluorescence, on the first day, anti-RECA-1 was added and the slides incubated overnight at 4°C conditions, then sections were incubated with the anti-mouse IgG labeled with Alexa Fluor 568 for 2 h. Thereafter, incubation with anti-VEGF-A and anti-GFAP antibodies was allowed overnight in the dark. Finally, anti-mouse IgG labeled with Alexa Fluor 488 and anti-rabbit IgG labeled with Alexa Fluor 647 were added for 2 h in the dark (Supplementary Table S1).

Negative control sections (no exposure to the primary antisera) were processed concurrently with the other sections for all immunohistochemical studies. Images were acquired using a motorized Leica DM6000 B microscope equipped with a DFC350FX camera (Leica, Mannheim, Germany).

The colocalization area was calculated using the “colocalization” plugin of ImageJ (after evaluating the threshold value for each channel) and expressed as percentage relative to the value of the VEGFR-1 or VEGF-A area. The mean fluorescence intensity of VEGF-A in control and oxaliplatin-treated animals was calculated by subtracting the background (multiplied by the total area) from the VEGF-A integrated intensity. Analyses were performed on three different images for each animal, collected through a 20x objective.

For confocal analysis, images were acquired with a Leica SP2 AOBS confocal microscope using a sequential scan setting (exciting lasers 488 nm and 561nm) to avoid channel bleed-through. Images were acquired though a 63x 1.4NA PL APO objective at voxel size of 232nm (xy) and 121nm (z). Confocal images were processed and analyzed using Fiji (Schindelin *et al*, 2012). Briefly, images were deconvolved using Deconvolution Lab2 with a synthetic PSF and ICTM algorithm (Sage *et al*, 2017). Colocalization analysis was performed with JACoP (Fiji plugin) (Bolte & Cordelières, 2006) and manually set thresholds. Colocalization parameters were calculated from 8 confocal z-stacks for each analysis and are given as mean ± SEM.

### Statistics

Results were expressed as means ± SEM and the analysis of variance was performed by ANOVA test. A Bonferroni’s significant difference procedure was used as post hoc comparison. P values less than 0.05 were considered significant. Data were analysed using “Origin® 10” software.

Electrophysiological data were analysed through one-way ANOVA followed by Dunnet’s multiple comparison post-hoc test for statistical significance within groups. Two-way ANOVA followed by Bonferroni post-hoc test for comparison between groups, by using GraphPad Prism 7.0.

## Acknowledgements

The authors are grateful to Prof. D. Salvemini (Department of Pharmacology and Physiology, Saint Louis University School of Medicine, St. Louis, MO, United States) and to Dr. Warren Glaab (Merck, USA) for their input in our work and for the editorial contribution to the manuscript.

## Competing Interests

The authors declare no potential conflicts of interest

## Financial Support

This work was supported by the IMI2 project NeuroDerisk (821528), the Italian Ministry of University and Research, the University of Florence, by the Italian Association for Cancer Research (AIRC) under IG 2017 - ID. 20353 project - PI Graziani Grazia, and by the Italian Ministry of Health RC18-2638151 to Pedro M. Lacal.

## Author contributions

LM performed treatments, behavioral measurements and drafted the manuscript, EL contributed to behavioral tests and analyzed data, CP, AP, AV and AT performed molecular assay, TM performed confocal analysis, SB, FR and SM performed electrophysiological measurements, GG, PML, SM, PF, AP and CG contributed to plan the study, drafted and revised the manuscript, LDCM conceived the study and drafted the manuscript.

## Notes

### Competing Interest Statement

The authors have declared no competing interest.

## References

Alles SRA & Smith PA (2018) Etiology and Pharmacology of Neuropathic Pain. Pharmacol Rev 70: 315–347

Argaw AT, Asp L, Zhang J, Navrazhina K, Pham T, Mariani JN, Mahase S, Dutta DJ, Seto J, Kramer EG, et al (2012) Astrocyte-derived VEGF-A drives blood-brain barrier disruption in CNS inflammatory disease. J Clin Invest 122: 2454–2468

Argyriou AA, Bruna J, Marmiroli P & Cavaletti G (2012) Chemotherapy-induced peripheral neurotoxicity (CIPN): An update. Crit Rev Oncol Hematol 82: 51–77

Atzori MG, Tentori L, Ruffini F, Ceci C, Bonanno E, Scimeca M, Lacal PM & Graziani G (2018) The Anti–Vascular Endothelial Growth Factor Receptor-1 Monoclonal Antibody D16F7 Inhibits Glioma Growth and Angiogenesis In Vivo. J Pharmacol Exp Ther 364: 77–86

Atzori MG, Tentori L, Ruffini F, Ceci C, Lisi L, Bonanno E, Scimeca M, Eskilsson E, Daubon T, Miletic H, et al (2017) The anti-vascular endothelial growth factor receptor-1 monoclonal antibody D16F7 inhibits invasiveness of human glioblastoma and glioblastoma stem cells. J Exp Clin Cancer Res 36: 106

Baptista-de-Souza D, Di Cesare Mannelli L, Zanardelli M, Micheli L, Nunes-de-Souza RL, Canto-de-Souza A & Ghelardini C (2014) Serotonergic modulation in neuropathy induced by oxaliplatin: effect on the 5HT2C receptor. Eur J Pharmacol 735: 141–149

Beazley-Long N, Hua J, Jehle T, Hulse RP, Dersch R, Lehrling C, Bevan H, Qiu Y, Lagrèze WA, Wynick D, et al (2013) VEGF-A165b Is an Endogenous Neuroprotective Splice Isoform of Vascular Endothelial Growth Factor A in Vivo and in Vitro. Am J Pathol 183: 918–929

Boccella S, Vacca V, Errico F, Marinelli S, Squillace M, Guida F, Di Maio A, Vitucci D, Palazzo E, De Novellis V, et al (2015) D-Aspartate Modulates Nociceptive-Specific Neuron Activity and Pain Threshold in Inflammatory and Neuropathic Pain Condition in Mice. Biomed Res Int 2015: 1–10

Bolte S & Cordelières FP (2006) A guided tour into subcellular colocalization analysis in light microscopy. J Microsc 224: 213–232

Branca JJV, Maresca M, Morucci G, Becatti M, Paternostro F, Gulisano M, Ghelardini C, Salvemini D, Di Cesare Mannelli L & Pacini A (2018) Oxaliplatin-induced blood brain barrier loosening: a new point of view on chemotherapy-induced neurotoxicity. Oncotarget 9: 23426– 23438

Cai M, Wang K, Murdoch CE, G. Y & Ahmed A (2017) Heterodimerisation between VEGFR-1 and VEGFR-2 and not the homodimers of VEGFR-1 inhibit VEGFR-2 activity. Vascul Pharmacol 88: 11–20

Cavaletti G, Tredici G, Petruccioli MG, Dondè E, Tredici P, Marmiroli P, Minoia C, Ronchi A, Bayssas M & Etienne GG (2001) Effects of different schedules of oxaliplatin treatment on the peripheral nervous system of the rat. Eur J Cancer 37: 2457–2463

Ceci C, Atzori MG, Lacal PM & Graziani G (2020) Role of VEGFs/VEGFR-1 Signaling and its Inhibition in Modulating Tumor Invasion: Experimental Evidence in Different Metastatic Cancer Models. Int J Mol Sci 21

Di Cesare Mannelli L, Lucarini E, Micheli L, Mosca I, Ambrosino P, Soldovieri MV, Martelli A, Testai L, Taglialatela M, Calderone V, et al (2017) Effects of natural and synthetic isothiocyanate-based H 2 S-releasers against chemotherapy-induced neuropathic pain: Role of Kv7 potassium channels. Neuropharmacology 121: 49–59

Di Cesare Mannelli L, Marcoli M, Micheli L, Zanardelli M, Maura G, Ghelardini C & Cervetto C (2015) Oxaliplatin evokes P2X7-dependent glutamate release in the cerebral cortex: A pain mechanism mediated by Pannexin 1. Neuropharmacology 97: 133–141

Di Cesare Mannelli L, Pacini A, Bonaccini L, Zanardelli M, Mello T & Ghelardini C (2013) Morphologic Features and Glial Activation in Rat Oxaliplatin-Dependent Neuropathic Pain. J Pain 14: 1585–1600

Di Cesare Mannelli L, Pacini A, Micheli L, Tani A, Zanardelli M & Ghelardini C (2014) Glial role in oxaliplatin-induced neuropathic pain. Exp Neurol 261: 22–33

Di Cesare Mannelli L, Tenci B, Micheli L, Vona A, Corti F, Zanardelli M, Lapucci A, Clemente AM, Failli P & Ghelardini C (2018) Adipose-derived stem cells decrease pain in a rat model of oxaliplatin-induced neuropathy: Role of VEGF-A modulation. Neuropharmacology 131: 166– 175

Chapouly C, Tadesse Argaw A, Horng S, Castro K, Zhang J, Asp L, Loo H, Laitman BM, Mariani JN, Straus Farber R, et al (2015) Astrocytic TYMP and VEGFA drive blood–brain barrier opening in inflammatory central nervous system lesions. Brain 138: 1548–1567

Christinger HW, Fuh G, de Vos AM & Wiesmann C (2004) The Crystal Structure of Placental Growth Factor in Complex with Domain 2 of Vascular Endothelial Growth Factor Receptor-1. J Biol Chem 279: 10382–10388

Cudmore MJ, Hewett PW, Ahmad S, Wang K-Q, Cai M, Al-Ani B, Fujisawa T, Ma B, Sissaoui S, Ramma W, et al (2012) The role of heterodimerization between VEGFR-1 and VEGFR-2 in the regulation of endothelial cell homeostasis. Nat Commun 3: 972

Davis-Smyth T, Presta LG & Ferrara N (1998) Mapping the Charged Residues in the Second Immunoglobulin-like Domain of the Vascular Endothelial Growth Factor/Placenta Growth Factor Receptor Flt-1 Required for Binding and Structural Stability. J Biol Chem 273: 3216– 3222

Failla C, Carbo M & Morea V (2018) Positive and Negative Regulation of Angiogenesis by Soluble Vascular Endothelial Growth Factor Receptor-1. Int J Mol Sci 19: 1306

Falcon BL, Chintharlapalli S, Uhlik MT & Pytowski B (2016) Antagonist antibodies to vascular endothelial growth factor receptor 2 (VEGFR-2) as anti-angiogenic agents. Pharmacol Ther 164: 204–225

Garraway SM & Huie JR (2016) Spinal Plasticity and Behavior: BDNF-Induced Neuromodulation in Uninjured and Injured Spinal Cord. Neural Plast 2016: 1–19

Ghelardini C, Galeotti N, Calvani M, Mosconi L, Nicolai R & Bartolini A (2002) Acetyl-l-carnitine induces muscarinic antinocieption in mice and rats. Neuropharmacology 43: 1180–1187

Graziani G, Ruffini F, Tentori L, Scimeca M, Dorio AS, Atzori MG, Failla CM, Morea V, Bonanno E, D’Atri S, et al (2016) Antitumor activity of a novel anti-vascular endothelial growth factor receptor-1 monoclonal antibody that does not interfere with ligand binding. Oncotarget 7: 72868–72885

Hamilton JL, Nagao M, Levine BR, Chen D, Olsen BR & Im H-J (2016) Targeting VEGF and Its Receptors for the Treatment of Osteoarthritis and Associated Pain: TARGETING VEGF AND VEGFRs FOR TREATMENT OF OSTEOARTHRITIS AND PAIN. J Bone Miner Res 31: 911–924

Hu X-M, Yang W, Du L-X, Cui W-Q, Mi W-L, Mao-Ying Q-L, Chu Y-X & Wang Y-Q (2019) Vascular Endothelial Growth Factor A Signaling Promotes Spinal Central Sensitization and Pain-related Behaviors in Female Rats with Bone Cancer. Anesthesiology 131: 1125–1147

Hulse RP, Beazley-Long N, Hua J, Kennedy H, Prager J, Bevan H, Qiu Y, Fernandes ES, Gammons M V, Ballmer-Hofer K, et al (2014) Regulation of alternative VEGF-A mRNA splicing is a therapeutic target for analgesia. Neurobiol Dis 71: 245–259

Hylden JLK & Wilcox GL (1980) Intrathecal morphine in mice: A new technique. Eur J Pharmacol 67: 313–316

Ibrahim EY & Ehrlich BE (2020) Prevention of chemotherapy-induced peripheral neuropathy: A review of recent findings. Crit Rev Oncol Hematol 145: 102831

Iyer S & Acharya KR (2011) Tying the knot: The cystine signature and molecular-recognition processes of the vascular endothelial growth factor family of angiogenic cytokines: Cystine-knot growth factors and angiogenesis. FEBS J 278: 4304–4322

Kolb NA, Smith AG, Singleton JR, Beck SL, Stoddard GJ, Brown S & Mooney K (2016) The Association of Chemotherapy-Induced Peripheral Neuropathy Symptoms and the Risk of Falling. JAMA Neurol 73: 860

Lacal, Pedro Miguel, Atzori, MG; Ruffini, F; Scimeca, M; Bonanno, E; Cicconi, R; Mattei, M; Bernardini, R; D’Atri, S; Tentori, L; Graziani, G (2020) Targeting the vascular endothelial growth factor receptor-1 by the monoclonal antobody D16F7 to increase the activity of immune checkpoint inhibitors against cutaneous melanoma. Pharmacol Res

Lacal PM & Graziani G (2018) Therapeutic implication of vascular endothelial growth factor receptor-1 (VEGFR-1) targeting in cancer cells and tumor microenvironment by competitive and non-competitive inhibitors. Pharmacol Res 136: 97–107

Lange C, Storkebaum E, de Almodóvar CR, Dewerchin M & Carmeliet P (2016) Vascular endothelial growth factor: a neurovascular target in neurological diseases. Nat Rev Neurol 12: 439–454

Lee GW, Son JY, Lee AR, Ju JS, Bae YC & Ahn DK (2019) Central VEGF-A pathway plays a key role in the development of trigeminal neuropathic pain in rats. Mol Pain 15: 174480691987260

Licht T & Keshet E (2013) Delineating multiple functions of VEGF-A in the adult brain. Cell Mol Life Sci 70: 1727–1737

Lin J, Li G, Den X, Xu C, Liu S, Gao Y, Liu H, Zhang J, Li X & Liang S (2010) VEGF and its receptor-2 involved in neuropathic pain transmission mediated by P2X2(/)3 receptor of primary sensory neurons. Brain Res Bull 83: 284–291

Lucarini E, Pagnotta E, Micheli L, Parisio C, Testai L, Martelli A, Calderone V, Matteo R, Lazzeri L, Di Cesare Mannelli L, et al (2019) Eruca sativa Meal against Diabetic Neuropathic Pain: An H(2)S-Mediated Effect of Glucoerucin. Molecules 24

McGaraughty S, Chu KL, Perner RJ, Didomenico S, Kort ME & Kym PR (2010) TRPA1 modulation of spontaneous and mechanically evoked firing of spinal neurons in uninjured, osteoarthritic, and inflamed rats. Mol Pain 6: 14

McGrath JC & Lilley E (2015) Implementing guidelines on reporting research using animals (ARRIVE etc.): new requirements for publication in BJP. Br J Pharmacol 172: 3189–3193

Meyer M (1999) A novel vascular endothelial growth factor encoded by Orf virus, VEGF-E, mediates angiogenesis via signalling through VEGFR-2 (KDR) but not VEGFR-1 (Flt-1) receptor tyrosine kinases. EMBO J 18: 363–374

Miltenburg NC & Boogerd W (2014) Chemotherapy-induced neuropathy: A comprehensive survey. Cancer Treat Rev 40: 872–882

Montague-Cardoso K, Pitcher T, Chisolm K, Salera G, Lindstrom E, Hewitt E, Solito E & Malcangio M (2020) Changes in vascular permeability in the spinal cord contribute to chemotherapy-induced neuropathic pain. Brain Behav Immun 83: 248–259

Nakayama M, Nakayama A, van Lessen M, Yamamoto H, Hoffmann S, Drexler HCA, Itoh N, Hirose T, Breier G, Vestweber D, et al (2013) Spatial regulation of VEGF receptor endocytosis in angiogenesis. Nat Cell Biol 15: 249–260

Nencini S, Ringuet M, Kim D-H, Greenhill C & Ivanusic JJ (2018) GDNF, Neurturin, and Artemin Activate and Sensitize Bone Afferent Neurons and Contribute to Inflammatory Bone Pain. J Neurosci 38: 4899–4911

Park JE, Chen HH, Winer J, Houck KA & Ferrara N (1994) Placenta growth factor. Potentiation of vascular endothelial growth factor bioactivity, in vitro and in vivo, and high affinity binding to Flt-1 but not to Flk-1/KDR. J Biol Chem 269: 25646–25654

Peach C, Mignone V, Arruda M, Alcobia D, Hill S, Kilpatrick L & Woolard J (2018) Molecular Pharmacology of VEGF-A Isoforms: Binding and Signalling at VEGFR2. Int J Mol Sci 19: 1264

Persico MG, Vincenti V & DiPalma T (1999) Structure, Expression and Receptor-Binding Properties of Placenta Growth Factor (PlGF). In Vascular Growth Factors and Angiogenesis, Claesson-Welsh L(ed) pp 31–40. Berlin, Heidelberg: Springer Berlin Heidelberg

Polomano RC, Mannes AJ, Clark US & Bennett GJ (2001) A painful peripheral neuropathy in the rat produced by the chemotherapeutic drug, paclitaxel: Pain 94: 293–304

Ponnambalam S & Alberghina M (2011) Evolution of the VEGF-Regulated Vascular Network from a Neural Guidance System. Mol Neurobiol 43: 192–206

Rojas DR, Tegeder I, Kuner R & Agarwal N (2018) Hypoxia-inducible factor 1α protects peripheral sensory neurons from diabetic peripheral neuropathy by suppressing accumulation of reactive oxygen species. J Mol Med 96: 1395–1405

Ruiz de Almodovar C, Lambrechts D, Mazzone M & Carmeliet P (2009) Role and Therapeutic Potential of VEGF in the Nervous System. Physiol Rev 89: 607–648

Russo R, D’Agostino G, Mattace Raso G, Avagliano C, Cristiano C, Meli R & Calignano A (2012) Central administration of oxytocin reduces hyperalgesia in mice: implication for cannabinoid and opioid systems. Peptides 38: 81–88

Sage D, Donati L, Soulez F, Fortun D, Schmit G, Seitz A, Guiet R, Vonesch C & Unser M (2017) DeconvolutionLab2: An open-source software for deconvolution microscopy. Methods 115: 28–41

Schindelin J, Arganda-Carreras I, Frise E, Kaynig V, Longair M, Pietzsch T, Preibisch S, Rueden C, Saalfeld S, Schmid B, et al (2012) Fiji: an open-source platform for biological-image analysis. Nat Methods 9: 676–682

Scholz J & Woolf CJ (2007) The neuropathic pain triad: neurons, immune cells and glia. Nat Neurosci 10: 1361–1368

Schratzberger P, Schratzberger G, Silver M, Curry C, Kearney M, Magner M, Alroy J, Adelman LS, Weinberg DH, Ropper AH, et al (2000) Favorable effect of VEGF gene transfer on ischemic peripheral neuropathy. Nat Med 6: 405–413

Seki T, Hosaka K, Fischer C, Lim S, Andersson P, Abe M, Iwamoto H, Gao Y, Wang X, Fong G-H, et al (2018) Ablation of endothelial VEGFR1 improves metabolic dysfunction by inducing adipose tissue browning. J Exp Med 215: 611–626

Selvaraj D, Gangadharan V, Michalski CW, Kurejova M, Stösser S, Srivastava K, Schweizerhof M, Waltenberger J, Ferrara N, Heppenstall P, et al (2015) A Functional Role for VEGFR1 Expressed in Peripheral Sensory Neurons in Cancer Pain. Cancer Cell 27: 780–796

Simone DA, Khasabov SG & Hamamoto DT (2008) Changes in response properties of nociceptive dorsal horn neurons in a murine model of cancer pain. Sheng Li Xue Bao 60: 635–644

Sommer C, Leinders M & Üçeyler N (2018) Inflammation in the pathophysiology of neuropathic pain: Pain 159: 595–602

Stevens M & Oltean S (2018) Modulation of VEGF-A Alternative Splicing as a Novel Treatment in Chronic Kidney Disease. Genes (Basel) 9: 98

Stockstill K, Doyle TM, Yan X, Chen Z, Janes K, Little JW, Braden K, Lauro F, Giancotti LA, Harada CM, et al (2018) Dysregulation of sphingolipid metabolism contributes to bortezomib-induced neuropathic pain. J Exp Med 215: 1301–1313

Taiana MM, Lombardi R, Porretta-Serapiglia C, Ciusani E, Oggioni N, Sassone J, Bianchi R & Lauria G (2014) Neutralization of Schwann Cell-Secreted VEGF Is Protective to In Vitro and In Vivo Experimental Diabetic Neuropathy. PLoS One 9: e108403

Telleria-Diaz A, Schmidt M, Kreusch S, Neubert A-K, Schache F, Vazquez E, Vanegas H, Schaible H-G & Ebersberger A (2010) Spinal antinociceptive effects of cyclooxygenase inhibition during inflammation: Involvement of prostaglandins and endocannabinoids: Pain 148: 26–35

Ved N, Da Vitoria Lobo ME, Bestall SM, L. Vidueira C Beazley-Long N, Ballmer-Hofer K, Hirashima M, Bates DO, Donaldson LF & Hulse RP (2018) Diabetes-induced microvascular complications at the level of the spinal cord: a contributing factor in diabetic neuropathic pain: Diabetes-induced microvascular degeneration and neuropathic pain. J Physiol 596: 3675–3693

Verheyen A, Peeraer E, Lambrechts D, Poesen K, Carmeliet P, Shibuya M, Pintelon I, Timmermans J-P, Nuydens R & Meert T (2013) Therapeutic potential of VEGF and VEGF-derived peptide in peripheral neuropathies. Neuroscience 244: 77–89

Verheyen A, Peeraer E, Nuydens R, Dhondt J, Poesen K, Pintelon I, Daniels A, Timmermans J-P, Meert T, Carmeliet P, et al (2012) Systemic anti-vascular endothelial growth factor therapies induce a painful sensory neuropathy. Brain 135: 2629–2641

Weng -RH, Cordella VJ & Dougherty MP (2003) Changes in sensory processing in the spinal dorsal horn accompany vincristine-induced hyperalgesia and allodynia: Pain 103: 131–138

Woolard J, Wang W-Y, Bevan HS, Qiu Y, Morbidelli L, Pritchard-Jones RO, Cui T-G, Sugiono M, Waine E, Perrin R, et al (2004) VEGF 165 b, an Inhibitory Vascular Endothelial Growth Factor Splice Variant: Mechanism of Action, In vivo Effect On Angiogenesis and Endogenous Protein Expression. Cancer Res 64: 7822–7835

