## Supplementary materials for "VEGF-A/VEGFR-1: a painful astrocyte-mediated signaling blocked by the anti-VEGFR-1 mAb D16F7"

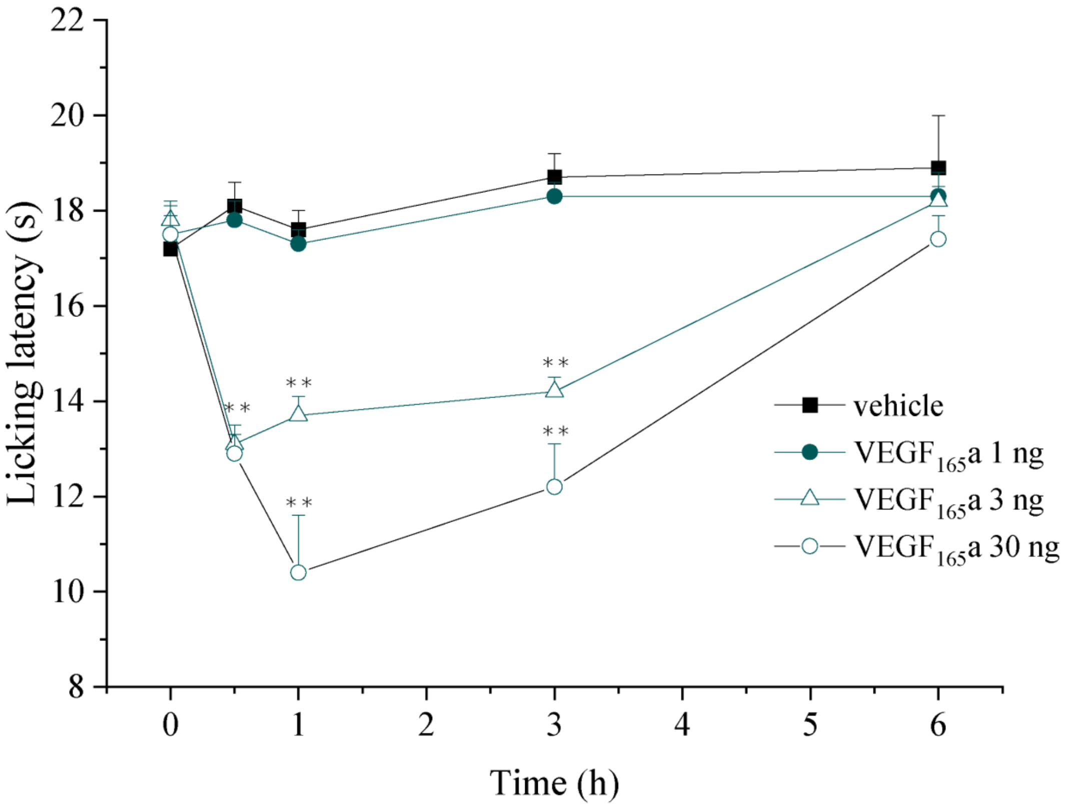


***Supplementary Figure S1. Nociceptive effect of VEGF$_{165}$a.*** *The pain threshold was measured by the Cold plate test over time after the intrathecal injection of VEGF$_{165}$a (n=5). Each value represents the mean ± SEM. **P<0.01 vs vehicle-treated animals. The analysis of variance was performed by One-way ANOVA. A Bonferroni's significant difference procedure was used as post hoc comparison.*

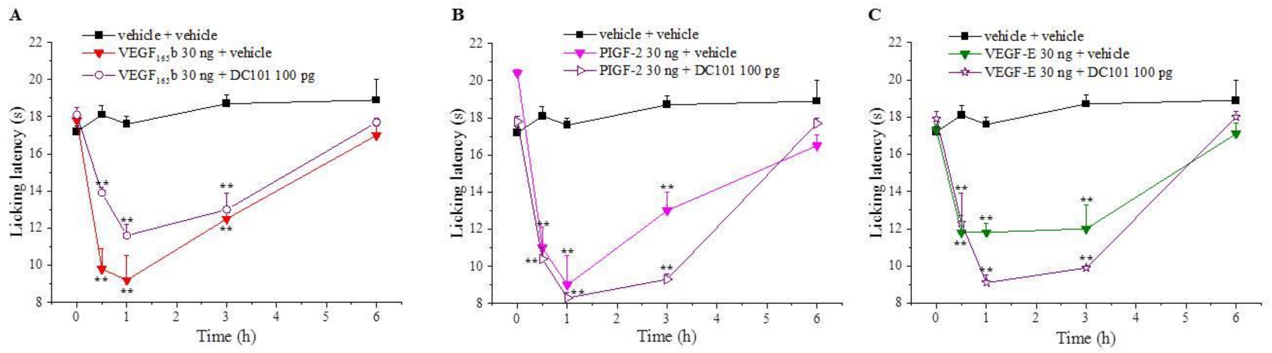

***Supplementary Figure S2. Hypersensitivity-induced by VEGF-A signalling modulators is not due to its interaction with VEGFR-2.*** *The response to a thermal stimulus (Cold plate test) was recorded after intrathecal infusion of (A) VEGF$_{165}$b 30 ng pretreated (15 min before) with DC101 (n=5), (B) PlGF-2 pretreated with DC101 (n=5), (C) VEGF-E pretreated with DC101 (n=5). Each value represents the mean ± SEM. **P<0.01 vs vehicle + vehicle-treated animals. The analysis of variance was performed by One-way ANOVA. A Bonferroni's significant difference procedure was used as post hoc comparison.*

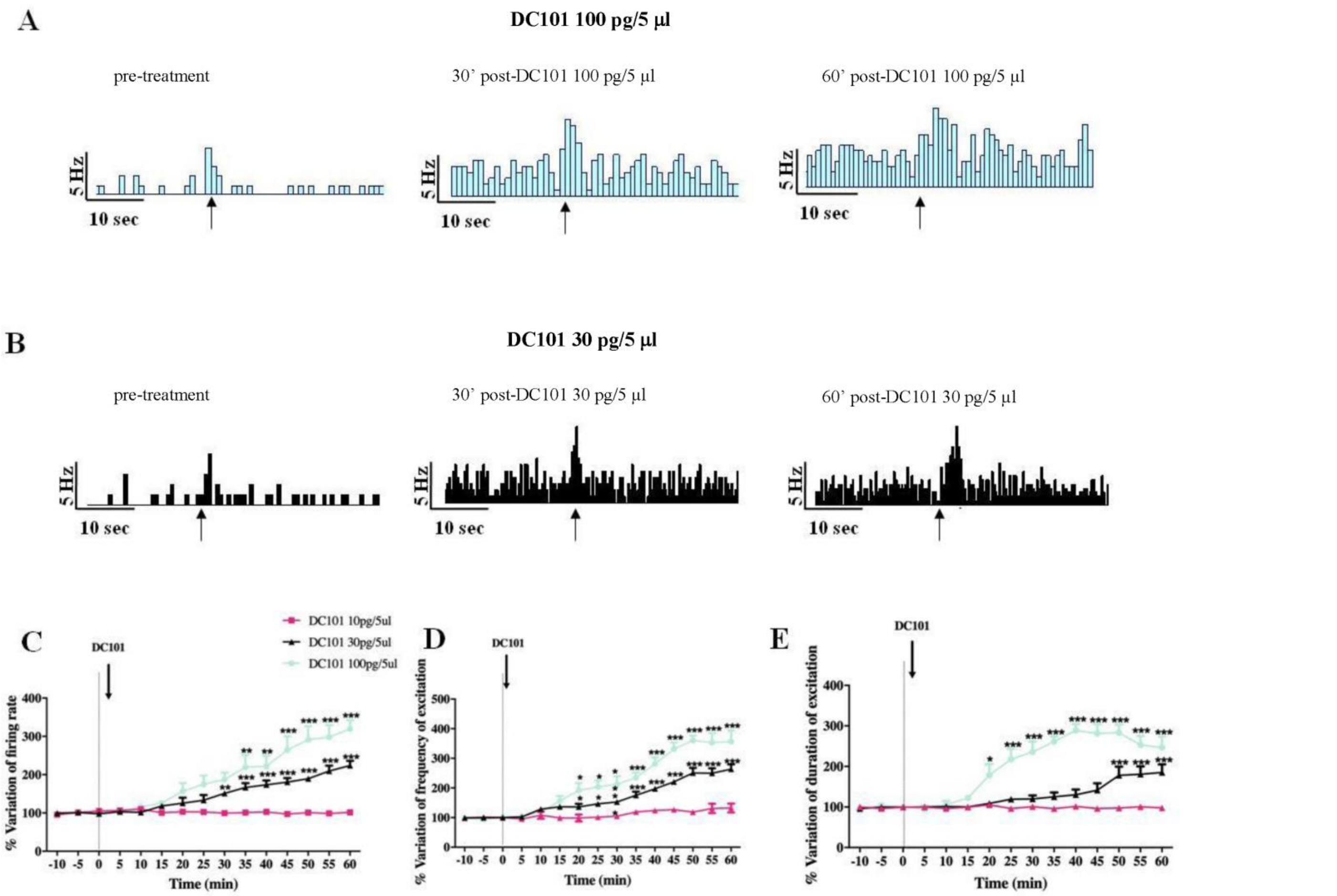


***Supplementary Figure S3. DC101 increases spontaneous and noxious-evoked activity of NS neurons.*** *Representative ratemeters showing spontaneous and noxious-evoked activity of NS neurons after spinal application of DC101 antibodies at 100 pg (A) and 30pg (B), black arrows indicate the noxious stimulation on the mouse hind-paw. Mean ± SEM population data of spinal cord application of DC101 (30 pg and 100 pg) on % variation of firing rate (C), % variation of frequency of excitation (D) and % variation of duration of evoked activity (E) of NS neurons in CD1 mice. Black arrows indicate vehicle, DC101 spinal application. Each point represents the mean of 5 different mice per group (one neuron recorded per each mouse). *P<0.05, **P<0.01 and ***P<0.001 indicate statistically difference vs pre-drug. One- way ANOVA followed by Dunnet's multiple comparison post-hoc test was performed for statistical significance within groups.*

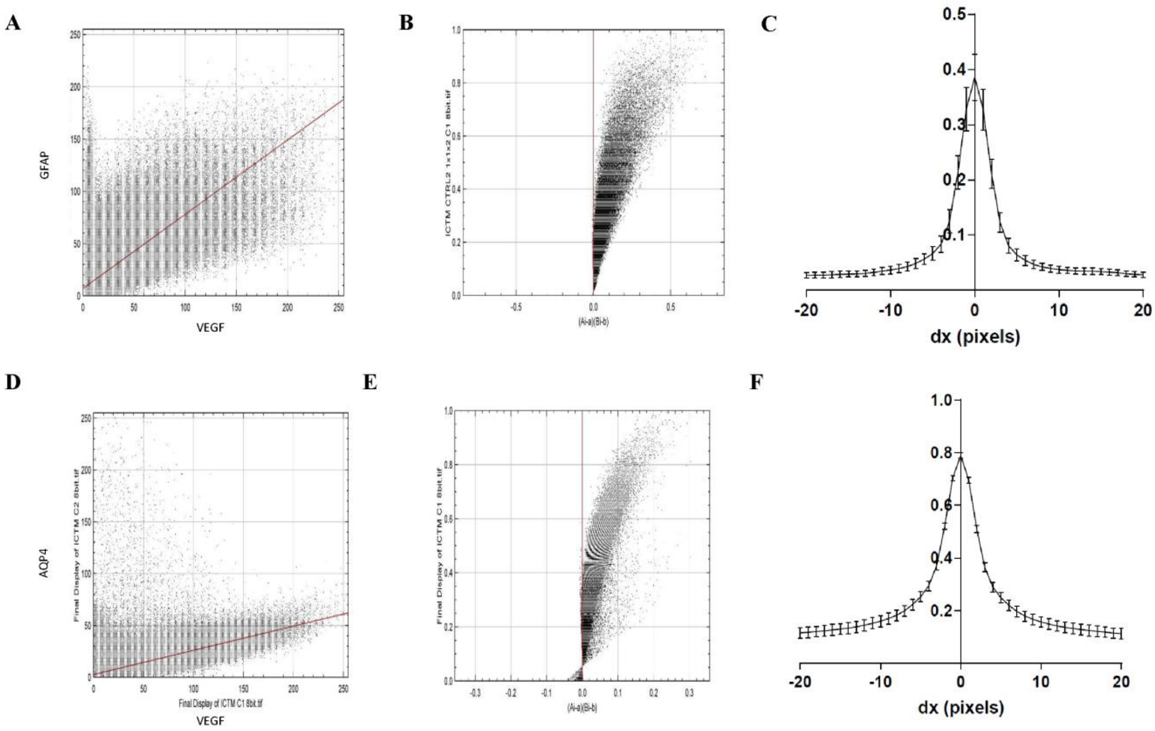


***Supplementary Figure S4. Analyses of GFAP and VEGF co-localization in confocal z-stacks.*** *A) Cytofluorogram relative to images in Fig. 5 B) Li's Intensity Correlation Analysis relative to images in Fig. 5. C) Van Steensel's Cross-Correlation Function (CCF), relative to all datasets (n=8, mean ± SEM). Pearson's correlation coefficient (PCC) is given by the CCF value corresponding to x =0. CCF at FWHM = 1.00 ± 0.04 μm (mean ± SEM, n=8). D, E)* ***Analyses ofAQP4 and VEGF co-localization in confocal z-stacks.*** *D) Cytofluorogram relative to images in Fig. 5 E) Li's Intensity Correlation Analysis relative to images in Fig. 5. CF Van Steensel's Cross-Correlation Function (CCF), relative to all datasets (n=8, mean ± SEM). Pearson's correlation coefficient (PCC) is given by the CCF value corresponding to x =0. CCF at FWHM =1.28±0.04 μm (mean ± SEM, n=8)*

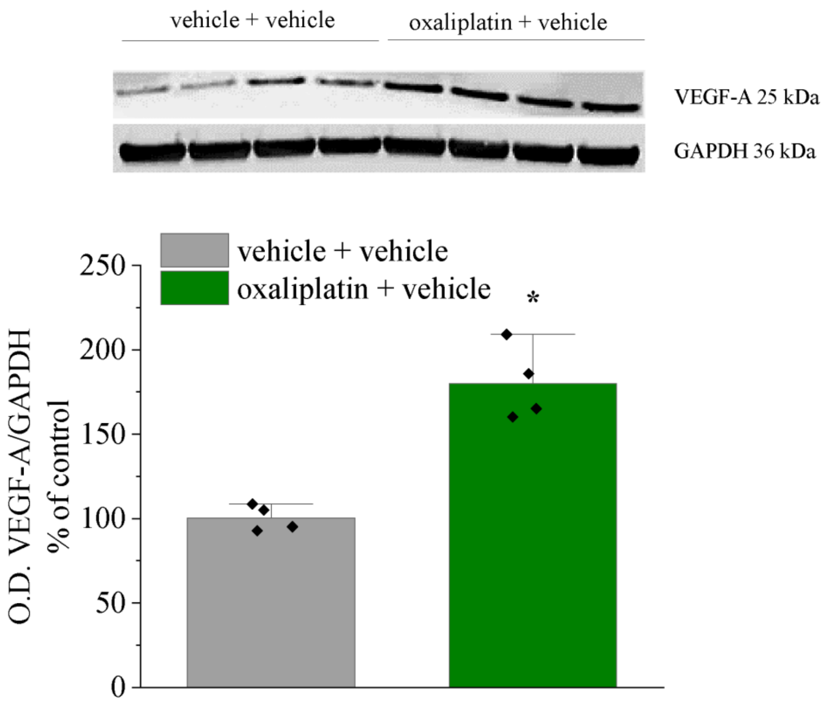


***Supplementary Figure S5. VEGF-A is increased in the spinal cord of mice with oxaliplatin-induced neuropathy.*** *Representative western blot images and densitometric analysis of VEGF-A expression in the lumbar section of the spinal cord of oxaliplatin-treated mice in comparison to control (n=4). Each value represents the mean ± SEM. *P<0.05 vs vehicle + vehicle group. The analysis of variance was performed by One-way ANOVA. A Bonferroni's significant difference procedure was used as post hoc comparison.*

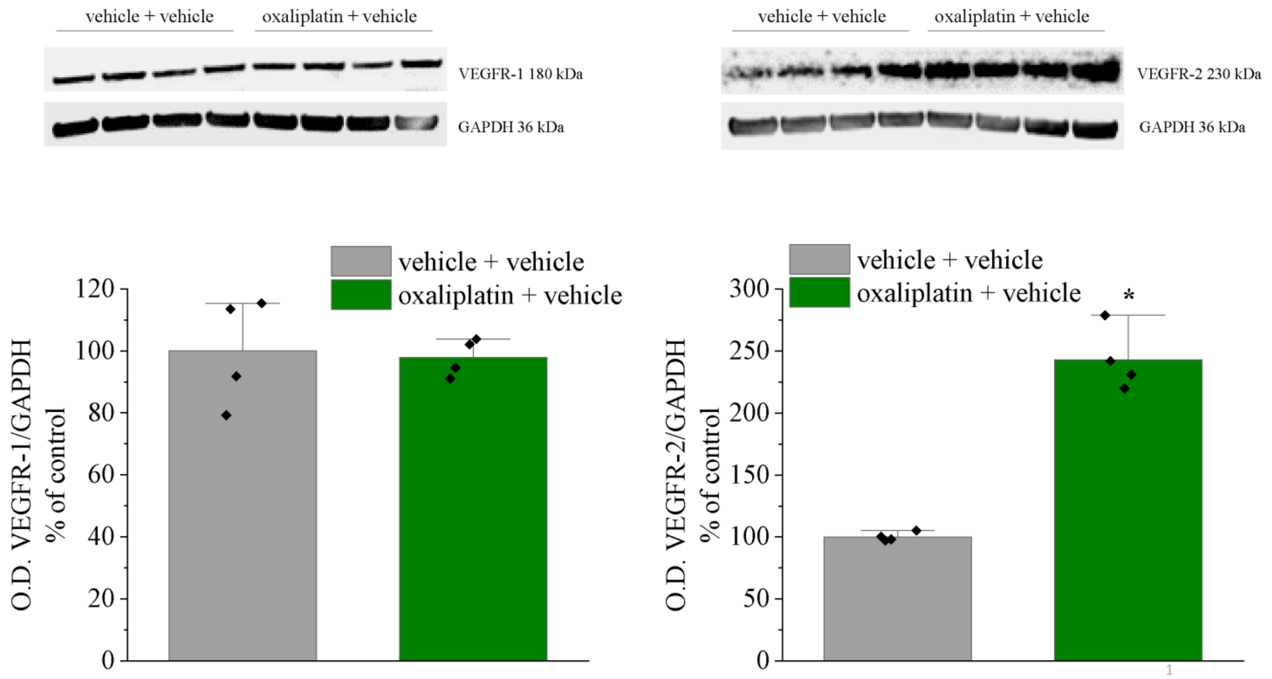


***Supplementary Figure S6. VEGFR-2 is increased in the spinal cord of mice with oxaliplatin-induced neuropathy.*** *Representative western blot images and densitometric analysis of VEGF-R1 and VEGFR-2 expression in the lumbar section of the spinal cord of oxaliplatin-treated mice in comparison to control (n=4). Each value represents the mean ± SEM. *P<0.05 vs vehicle + vehicle group. The analysis of variance was performed by One-way ANOVA. A Bonferroni's significant difference procedure was used as post hoc comparison.*

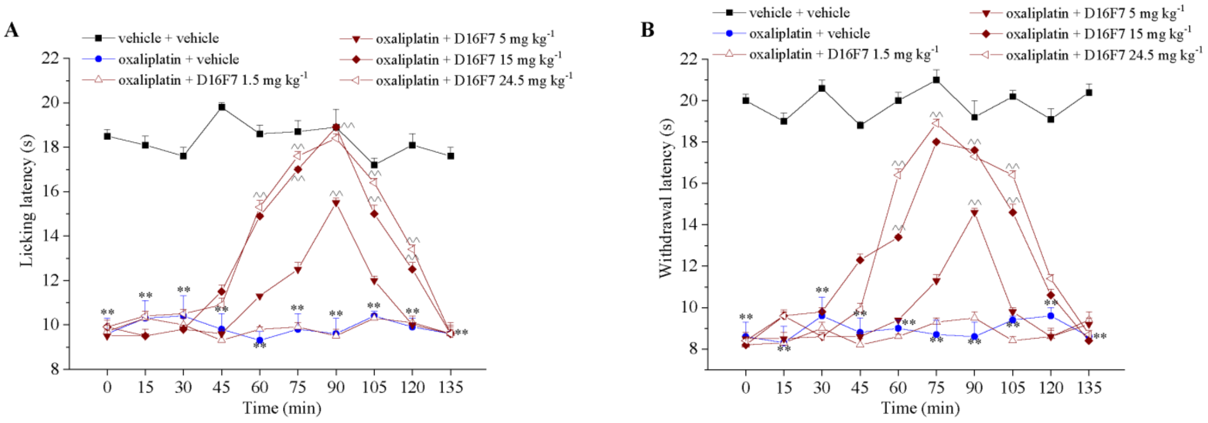


***Supplementary Figure S7. D16F7 mAb reduced oxaliplatin-induced pain after systemic administration.*** *Effect of D16F7 mAb evaluated by (A) Cold plate and (B) Paw pressure tests in a mouse model of oxaliplatin-induced neuropathy after i.p. injection (A, B, n=6). Each value represents the mean ± SEM. **P<0.01 vs vehicle + vehicle-treated animals; ^^P<0.01 vs oxaliplatin + vehicle-treated animals. The analysis of variance was performed by One-way ANOVA. A Bonferroni's significant difference procedure was used as post hoc comparison.*

**Supplementary Table S1. List of antibodies used for immunohistochemistry and western blot assays**

| Target | Antigen | Supplier | Catalog# | Antibody | Host | Usage | Conc. | Analysis |
|---|---|---|---|---|---|---|---|---|
| Astrocytes | GFAP | Merck Millipore | MAB3402X | Monoclonal conj. 488 | Ms | Primary | 1:500 | IF |
| Astrocytes | GFAP | Dako | ZO334 | Polyclonal | Rb | Primary | 1:500 | IF |
| Neurons | NeuN | Merck Millipore | MAB377X | Monoclonal conj. 488 | Ms | Primary | 1:500 | IF |
| Microglia | Iba-1 | Wako | 016-20001 | Polyclonal | Rb | Primary | 1:200 | IF |
| Vascular Endothelial Growth Factor Receptor-1 | VEGFR1 | Bioss | bs-0170R | Polyclonal | Rb | Primary | 1:100 | IF |
| Vascular Endothelial Growth Factor-A | VEGF-A | Santa Cruz Biotechnology | sc-7269 | Monoclonal | Ms | Primary | 1:100 | IF |
| Vascular Endothelial Growth Factor | VEGF | BD Pharmigen | 555036 | Monoclonal | Ms | Primary | 1:1000 | WB |
| Vascular Endothelial Growth Factor Receptor-1 | VEGFR-1 | Abcam | 32152 | Monoclonal | Rb | Primary | 1:1000 | WB |
| Vascular Endothelial Growth Factor Receptor-2 | VEGFR2/ DC101 | Bio X Cell | BE0060 | Monoclonal | Ms | Primary | 1:5000 | WB |
| Aquaporin 4 | AQP-4 | Santa Cruz Biotechnology | sc-32739 | Monoclonal | Ms | Primary | 1:100 | IF |
| RECA-1 | RECA-1 | Santa Cruz Biotechnology | Sc-52665 | Monoclonal | Ms | Primary | 1:100 | IF |
| Glyceraldehyde 3-phosphate dehydrogenase | GAPDH | Santa Cruz Biotechnology | sc-32233 | Monoclonal | Ms | Primary | 1:2500 | WB |
| Rabbit FC | Rabbit FC | Life technologies | A-11011 | Polyclonal | Rb | Secondary Alexa Fluor 568 | 1:500 | IF |
| Mouse FC | Mouse FC | Life technologies | A-11004 | Polyclonal | Ms | Secondary Alexa Fluor 568 | 1:500 | IF |
| Mouse FC | Mouse FC | Life technologies | A-11001 | Polyclonal | Ms | Secondary Alexa Fluor 488 | 1:500 | IF |
| Rabbit FC | Rabbit FC | Life technologies | A-21443 | Polyclonal | Rb | Secondary Alexa Fluor 647 | 1:200 | IF |
| 4’, 6-diamidin-2-fenilindolo | DAPI | Thermo scientific | 62248 | N.A. | N.A. | N.A. | 1:2000 | IF |

| | | | | | | | | |
|---|---|---|---|---|---|---|---|---|
| Rabbit-HRP | r-IgG-h | Bethyl | A120-201P | Polyclonal | Rb | Secondary conj. HRP | 1:5000 | WB |
| Mouse-HRP | m-IgGk BP-HRP | Santa Cruz Biotechnology | Sc-516102 | N.A. | Ms | Secondary conj. HRP | 1:5000 | WB |
| α-tubulin | α-4a | Sigma-Aldrich | T6074 | Monoclonal | Ms | Secondary | 1:5000 | WB |

**Supplementary Table S2. Hole Board test**

| | Day 3 | | Day 5 | | Day 9 | |
|---|---|---|---|---|---|---|
| Treatments | hole | board | hole | board | hole | board |
| vehicle + vehicle | 46.8 ± 10.1 | 71.2 ± 11.1 | 27.4 ± 5.2 | 56.5 ± 6.0 | 21.0 ± 1.0 | 37.0 ± 1.5 |
| vehicle + oxaliplatin | 57.5 ± 4.1 | 66.4 ± 6.6 | 33. 8 ± 7.5 | 44.6 ± 5.8 | 18.0 ± 1.4 | 39.2 ± 4.0 |
| scrambled + oxaliplatin | 49.2 ± 6.0 | 64.3 ± 8.5 | 25.9 ± 6.3 | 47.3 ± 3.8 | 24.4 ± 3.6 | 36.8 ± 2.6 |
| VEGFA-shRNAmir + oxaliplatin | 65.8 ± 4.8 | 174.4 ± 18.3** | 38.5 ± 6.6 | 69.6 ± 6.7 | 22.8 ± 2.7 | 40.4 ± 3.6 |

The Hole board test was performed 3, 5 and 9 days after the beginning of oxaliplatin treatment (n=5). Each value represents the mean ± SEM. **P<0.01 vs vehicle + vehicle treated animals. The analysis of variance was performed by One-way ANOVA. A Bonferroni's significant procedure was used as post hoc comparison.
